# Qualifying a human Liver-Chip for predictive toxicology: Performance assessment and economic implications

**DOI:** 10.1101/2021.12.14.472674

**Authors:** Lorna Ewart, Athanasia Apostolou, Skyler A. Briggs, Christopher V. Carman, Jake T. Chaff, Anthony R. Heng, Sushma Jadalannagari, Jeshina Janardhanan, Kyung-Jin Jang, Sannidhi R. Joshipura, Mahika M. Kadam, Marianne Kanellias, Ville J. Kujala, Gauri Kulkarni, Christopher Y. Le, Carolina Lucchesi, Dimitris V. Manatakis, Kairav K. Maniar, Meaghan E. Quinn, Joseph S. Ravan, Ann Catherine Rizos, John F.K. Sauld, Josiah D. Sliz, William Tien-Street, Dennis Ramos Trinidad, James Velez, Max Wendell, Onyi Irrechukwu, Prathap Kumar Mahalingaiah, Donald E. Ingber, Jack W. Scannell, Daniel Levner

## Abstract

Human organ-on-a-chip (Organ-Chip) technology has the potential to disrupt preclinical drug discovery and improve success in drug development pipelines as it can recapitulate organ-level pathophysiology and clinical responses. The Innovation and Quality (IQ) consortium formed by multiple pharmaceutical and biotechnology companies to confront this challenge has published guidelines that define criteria for qualifying preclinical models, however, systematic and quantitative evaluation of the predictive value of Organ-Chips has not yet been reported. Here, 870 Liver-Chips were analyzed to determine their ability to predict drug-induced liver injury (DILI) caused by small molecules identified as benchmarks by the IQ consortium. The Liver-Chip met the qualification guidelines across a blinded set of 27 known hepatotoxic and non-toxic drugs with a sensitivity of 87% and a specificity of 100%. A computational economic value analysis suggests that with this performance the Liver-Chip could generate $3 billion annually for the pharmaceutical industry due to increased R&D productivity.

## Introduction

Despite billion-dollar investments in research and development, the process of approving new drugs remains lengthy and costly due to high attrition rates^1,2,3^. Failure is common because the models used preclinically—which include computational, traditional cell culture, and animal models—have limited predictive validity^4^. The resulting damage to productivity in the pharmaceutical industry causes concern across a broad community of drug developers, investors, payers, regulators, and patients, the last of whom desperately need access to medicines with proven efficacy and improved safety profiles. Approximately 75% of the cost in research and development is the cost of failure^5^— that is, money spent on projects in which the candidate drug was deemed efficacious and safe by early testing but was later revealed to be ineffective or unsafe in human clinical trials. Pharmaceutical companies are addressing this challenge by learning from drugs that failed and devising frameworks to unite research and development organizations to enhance the probability of clinical success^6,7,8,9^. One of the major goals of this effort is to develop preclinical models that could enable a “fail early, fail fast” approach, which would result in candidate drugs with greater probability of clinical success, improved patient safety, lower cost, and a faster time to market.

There are significant practical challenges in ascertaining the predictive validity of new preclinical models, as there is a broad diversity of chemistries and mechanisms of action or toxicity to consider, as well as significant time needed to confirm the model’s predictions once tested in the clinic. Consequently, arguments for the adoption of these new models are often based on features that are *presumed* to correlate with human responses to pharmacological interventions— realistic histology, similar genetics, or the use of patient-derived tissues. But even here there is a common problem in much of the academic literature: the important model features are chosen *post-hoc* by the authors and not prospectively by an independent third party that has expertise in the therapeutic problem at hand^10^.

The Innovation and Quality (IQ) consortium is a collaboration of pharmaceutical and biotechnology companies that aims to advance science and technology to enhance drug discovery programs. To further this goal, the consortium has described a series of performance criteria that a new preclinical model must meet to become qualified. Within this consortium is an affiliate dedicated to microphysiological systems (MPS), which include organ-on-a-chip (Organ-Chip) technology that employs microfluidic engineering to recapitulate *in vivo* cell and tissue microenvironments in an organ-specific context^11,12^. This is achieved by recreating tissue-tissue interfaces and providing fine control over fluid flow and mechanical forces^13,14^, optionally including supporting interactions with immune cells^15^ and microbiome^16^, and reproducing clinical drug exposure profiles^17^. Recognizing the promise of MPS for drug research and development, the IQ MPS affiliate has provided guidelines for qualifying new models for specific contexts of use to help advance regulatory acceptance and broader industrial adoption^18^; however, to this date, there have been no publications describing studies that carry out this type of performance validation for any specific context of use or that demonstrate an MPS capable of meeting these IQ consortium performance goals.

Guided by the IQ MPS affiliate’s roadmap on liver MPS^19^, which states that *in vitro* models for predicting drug-induced liver injury (DILI) that meet its guidelines are more likely to exhibit higher predictive validity than those that do not, we rigorously assessed commercially available human Liver-Chips (from Emulate, Inc.) within the context of use of DILI prediction. In this study, we tested 870 Liver-Chips using a blinded set of 27 different drugs with known hepatotoxic or non-toxic behavior recommended by the IQ consortium (Table 1). We compared the results to the historical performance of animal models as well as 3D spheroid cultures of primary human hepatocytes, which are preclinical models that are frequently employed in this context of use in the pharmaceutical industry^20^. In addition, we analyzed the Liver-Chip results from an economic perspective by estimating the financial value they could offer through their use in preclinical development in supporting toxicity-related decisions. We conclude with recommendations on how this type of platform might be implemented in pharmaceutical industry screening programs.

**Table 1.** Small Molecule Drug Compounds used in the Liver-Chip evaluation. The 27 small molecule drugs are listed according to the IQ MPS affiliate classification and their ranking in the Garside DILI severity category, where 1 corresponds to drugs with severe clinical DILI and 5 to those with no DILI26,42. Structurally related toxic and non-toxic pairs are indicated as well using bold, italic text.

| Drug | IQ MPS List | Tested in Spheroid | Spheroid False Negative | Garside DILI Rank |
| --- | --- | --- | --- | --- |
| Ambrisentan | ***Yes, matched with Sitaxsentan*** | Yes | No | 5 |
| Asunaprevir | Yes, no matched pair | No | No | 2 |
| Benoxaprofen | No | Yes | Yes | 1 |
| Beta-Estradiol | No | Yes | Yes | 3 |
| Buspirone | ***Yes, matched with Nefazodone*** | Yes | No | 4 |
| Chlorpheniramine | No | Yes | Yes | 3 |
| Clozapine | ***Yes, matched with Olanzapine*** | Yes | No | 2 |
| Diclofenac | Yes, no matched pair | Yes | No | 2 |
| Entacapone | ***Yes, matched with Tolcapone*** | Yes | No | 4 |
| Fialuridine | ***Yes, matched with FIRU*** | Yes | No | 1 |
| FIRU | ***Yes, matched with Fialuridine*** | No | No | 5 |
| Labetalol | No | Yes | Yes | 1 |
| Levofloxacin | ***Yes, matched with Trovafloxacin*** | Yes | Yes | 2 |
| Lomitapide | No, Mipomersen substitute | No | No | 3 |
| Nefazodone | ***Yes, matched with Buspirone*** | Yes | No | 1 |
| Olanzapine | ***Yes, matched with Clozapine*** | No | No | 5 |
| Pioglitazone | ***Yes, matched with Troglitazone*** | Yes | Yes | 3 |
| Simvastatin | No | Yes | Yes | 2 |
| Sitaxsentan | ***Yes, matched with Ambrisentan*** | Yes | No | 1 |
| Stavudine | No | Yes | Yes | 1 |
| Tacrine | No | Yes | Yes | 2 |
| Telithromycin | Yes, no matched pair | No | No | 1 |
| Tolcapone | ***Yes, matched with Entacapone*** | Yes | No | 1 |
| Troglitazone | ***Yes, matched with Pioglitazone*** | Yes | No | 1 |
| Trovafloxacin | ***Yes, matched with Levofloxacin*** | Yes | No | 1 |
| Ximelagatran | No | Yes | Yes | 1 |
| Zileuton | Yes, no matched pair | Yes | Yes | 2 |

## Results

### Liver-Chip satisfies IQ MPS affiliate guidelines

The IQ guidelines for assessment of an *in vitro* liver MPS within the DILI prediction context of use requires evidence that the model replicates key histological structures and functions of the liver; furthermore, the model must be able to distinguish between 7 pairs of small molecule toxic drugs and their non-toxic structural analogs. If the model passes through these hurdles, it must demonstrate its ability to predict the clinical responses of 6 additional selected drugs.

The Liver-Chips that we evaluated against these standards contain two parallel microfluidic channels separated by a porous membrane. Following the manufacturer’s instructions, primary human hepatocytes are cultured between two layers of extracellular matrix (ECM) in the upper ‘parenchymal’ channel, while primary human liver sinusoidal endothelial cells (LSECs), Kupffer cells, and stellate cells are placed in the lower ‘vascular’ channel in ratios that approximate those observed *in vivo* (Figure 1). All cells passed quality control criteria that included post-thaw viability >90%, low passage number (preferably P3 or less), and expression of cell-specific markers. Similar results were obtained using hepatocytes from three different human donors, which were procured from the same commercial vendor (Supplementary Table S1).

**Figure 1.**
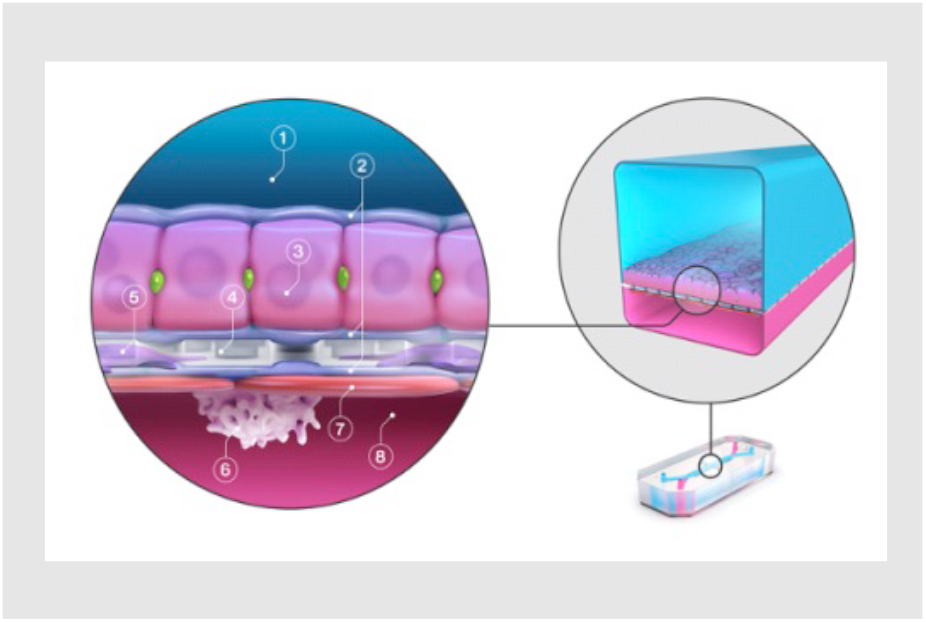
Schematic of the Liver-Chip showing primary human hepatocytes (3) that are sandwiched within an extracellular matrix (2) on a porous membrane (4) within the upper parenchymal channel (1), while human liver sinusoidal endothelial cells (7), Kupffer cells (6), and stellate cells (5) are cultured on the opposite side of the membrane in the lower vascular channel (8).

Live microscopy of the Liver-Chips revealed a continuous monolayer of hepatocytes displaying cuboidal and binucleated morphology in the upper ‘parenchymal’ channel of the chips, as well as a monolayer of polygonal shaped LSECs in the bottom ‘vascular’ channel, on the opposite side of the porous membrane (Figure 2a). Confocal fluorescence microscopy also confirmed liver-specific morphological structures as indicated by the presence of differentiation markers, including bile canaliculi containing a polarized distribution of F-actin and multidrug resistance-associated protein 2 (MRP2; Figure 2b), hepatocytes rich with mitochondrial membrane ATP synthase beta subunit (ATPB; Figure 2b), PECAM-1 (CD31) expressing LSECs, CD68^+^ Kupffer cells, and desmin-containing stellate cells (Figure 2c). In addition, transmission electron microscopy confirmed the existence of similar cell-cell relationships and structures to those found in human liver, including well developed junction-lined bile canaliculi and adhesions between Kupffer cells and sinusoidal endothelial cells (Supplementary Figure S1).

**Figure 2.**
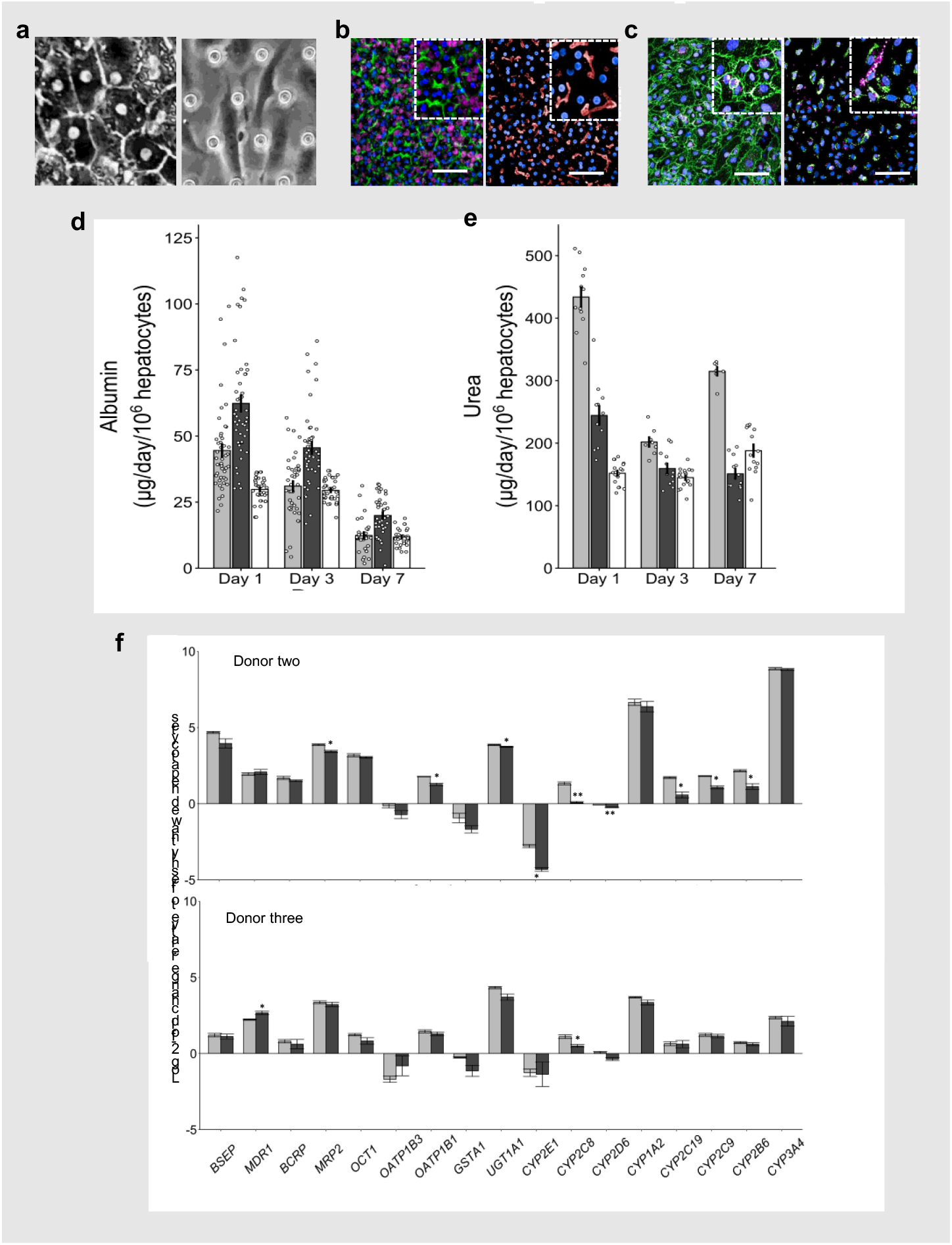
Recapitulation of human liver structure and function in the Liver-Chip. **a)** Representative phase contrast microscopic images of hepatocytes in the upper channel of Liver-Chip (left) and non-parenchymal cells in the lower vascular channel (right); the regular array of circles are the pores in the membrane in the right image. **b)** Representative immunofluorescence microscopic images showing the phalloidin stained actin cytoskeleton (green) and ATPB containing mitochondria (magenta) (left), MRP2-containing bile canaliculi (red) (right). **c)** CD31-stained liver sinusoidal endothelial cells (green) and desmin-containing stellate cells (magenta) (left), and CD68+ Kupffer cells (green; right) co-localized with desmin-containing stellate cells (magenta; right). All images in b) and c) show DAPI-stained nuclei (blue) (bar, 100 µm; inset is shown at 5 times higher magnification). **d)** Albumin and **e)** urea levels in the effluent from the upper channels of vehicle-treated Liver-Chips created with cells from 3 different donors (light and dark grey bars represent donor one and two respectively, white bars represent donor three) on days 1, 3 and 7 post-vehicle administration, measured by ELISA. Data are presented as mean ± standard error of the mean (S.E.M.) with N = 29 to 46 and N = 7 to 18 chips for albumin and urea respectively across all days and donors. **f)** Levels of key liver-specific genes in control Liver-Chips as determined by RNA-seq analysis on days 3 (light grey) and 7 (dark grey) post-vehicle administration with donor two in the upper panel and donor three in the lower panel. Data are presented as Log2 (fold change) of the TPM (Transcript Per Million) expression relative to freshly thawed hepatocytes with N = 4 chips; statistical significance of values between day 3 and 7 was determined using a paired t-test; *, p<0.05, **, p <0.01.

Albumin and urea production are widely accepted as functional markers for cultured hepatocytes with the goal of reaching computed production levels observed in human liver *in vivo* (∼ 20-105 µg and 56-159 µg per 10^6^ hepatocytes per day, respectively)^19,21^. Liver-Chips fabricated with cells from three different hepatocyte donors were able to maintain physiologically relevant levels of albumin and urea synthesis over 1 week in culture (Figure 2d and 2e). Importantly, in line with the IQ MPS guidelines, the coefficient of variation for the mean daily production rate of urea was always below 5% in all donors on day 1 but increased to 20% on day 7; however, it was higher for albumin production across all donors on day 1 but was between 14 to 27% by day 7. These data corroborate the reproducibility and robustness of the Liver-Chip across experiments and highlight variability across donors that is not unlike the variability observed in humans. In fact, it is important to be able to analyze and understand donor-to-donor variability when evaluating cell-based platforms for the prediction of clinical outcome^22^ or when a drug moves into clinical studies.

Because hepatocytes maintained in conventional static cultures rapidly reduce transcription of relevant liver-specific genes^23^, the IQ MPS guidelines require confirmation that the genes representing major Phase I and II metabolizing enzymes, as well as uptake and efflux drug transporters, are expressed and that their levels of expression are stable. On days 3 and 7 post-vehicle administration, compared to freshly thawed hepatocytes, we detected high levels of expression in both donors for 13 of the 17 genes requested by IQ MPS, confirming that the chip provides a suitable microenvironment to maintain hepatocytes. Gene expression was significantly lower on day 7 compared to day 3 for CYP2D6, CYP2C8, CYP2E1 and MRP2 in donor two and only for MRP2 in donor 3 (Figure 2f). Gene expression levels were lower than freshly thawed hepatocytes for genes encoding OATP1B3, GSTA1, CYP2E1 and CYP2D6, a profile reflected in two donors. Moreover, the demonstration that CY2C9 and CYP3A4 gene expression is maintained above freshly thawed hepatocytes for 7 days post-vehicle or drug administration is encouraging as together the CYP2C and CYP3A families make up 50% of the total CYP population^75^. CYP3A4 is also the major enzyme that metabolizes many marketed drugs. Previously, the same Liver-Chip has been shown to exhibit Phase I and II functional activities that are comparable to freshly isolated human hepatocytes and 3D hepatic spheroids^21,25^ as well as superior activity relative to hepatocytes in a 2D sandwich-assay plate configuration^21^. Taken together, these data support the notion that the Liver-Chip provides a good microenvironment for hepatocytes to maintain functionality.

As these data confirmed that the Liver-Chip meets the major structural characterization and basic functionality requirements stipulated by the IQ MPS guidelines, we then carried out studies to evaluate this human model as a tool for DILI prediction. IQ MPS identified 7 pairs of small-molecule drugs where one drug has been reported to produce DILI in clinical studies and their structural analog was inactive or exhibited a lower activity and did not produce clinical DILI (Table 1). Past work in the MPS field has focused on technically accessible endpoints that can be easily measured but are unfortunately not clinically relevant or translatable (e.g., IC_50_ forreduction in total ATP content)^26,27^. Furthermore, although cytotoxicity measures are fundamental in the assessment of a drug’s potential for hepatotoxicity *in vitro* ^28,29^, gene expression and various phenotypic changes can occur at much lower concentrations^30,31^. As the Liver-Chip enables multiple measures of drug effects and use of multiple measures may provide further sensitivity and add value^32^, we assessed drug toxicities on days 1, 3 and 7 post-drug or vehicle administration by quantifying both inhibition of albumin production as a general measure of hepatocellular functionality and increases in release of alanine aminotransferase (ALT) protein, which is used clinically as a measure of liver damage. We also scored hepatocyte injury using morphological analysis at 1, 3, and 7 days after drug or vehicle exposure, where higher injury scores indicated greater cellular injury.

We tested the 7 toxic drugs across 8 concentrations that bracket the human plasma C_max_ for each drug based on free (non-protein bound) drug concentrations, with the highest concentrations at 300x C_max_ (unless not permitted by solubility limits as was found for levofloxacin) to represent clinically relevant test concentrations for *in vitro* models^33^ (Supplementary Table S2). The known toxic compounds showed clear concentration- and time-dependent patterns that varied depending on compound. Typically, when albumin production was inhibited, morphological injury scores and ALT levels also increased, but we found that a decrease of albumin production was the most sensitive marker of hepatocyte toxicity in the Liver-Chip, as shown in sample paired comparisons of clozapine and olanzapine, troglitazone and pioglitazone, and trovafloxacin and levofloxacin (Figure 3a). Importantly, all 7 of the toxic drugs reduced albumin production or resulted in an increase in ALT protein or injury morphology scores at lower multiples of the free human C_max_ compared to each of their non-toxic comparators, a finding that was repeated across 3 donors (Table 2). Furthermore, immunofluorescence microscopic imaging for markers of apoptotic cell death (caspase 3/7) and mitochondrial injury measured by visualizing reduction of tetramethylrhodamine methyl ester (TMRM) accumulation (Figure 3b) provided confirmation of toxicity and, in many cases, provided some insight into the potential mechanism of toxicity. For example, the third-generation anti-infective trovafloxacin is believed to have an inflammatory component to its toxicity, potentially mediated by Kupffer cells, but this is only seen in animal models if an inflammatory stimulant such as lipopolysaccharide (LPS) is co-administered^34^. Interestingly, immunofluorescence microscopic imaging of the Liver-Chip revealed that there was a concentration-dependent increase in caspase 3/7 staining following trovafloxacin treatment (Figure 3b, top panel); this supports a potential apoptotic component to its toxicity. Of note, levofloxacin, the lesser toxic structural analog, did not cause cellular apoptosis and trovafloxacin caused it without an inflammatory stimulant. The role of an activated immune system is considered to contribute to idiosyncratic DILI where a reactive metabolite forms an adduct that behaves like a hapten to activate the adaptive immune system^77^ or directly activates innate immune cells (e.g., Kupffer cell) to increase inflammatory cytokine production such as TNFa^76^. To assess whether trovafloxacin was able to activate Kupffer cells in the absence of an inflammatory stimulant we measured IL-6 and TNFa in top channel effluent of Liver-Chips from each of the three donors. We were unable to see any concentration-dependent increase in cytokine production (data not shown) suggesting that an additional inflammatory stimulus may be required, which in turn will likely exacerbate the toxicity of trovafloxacin.

**Figure 3.**
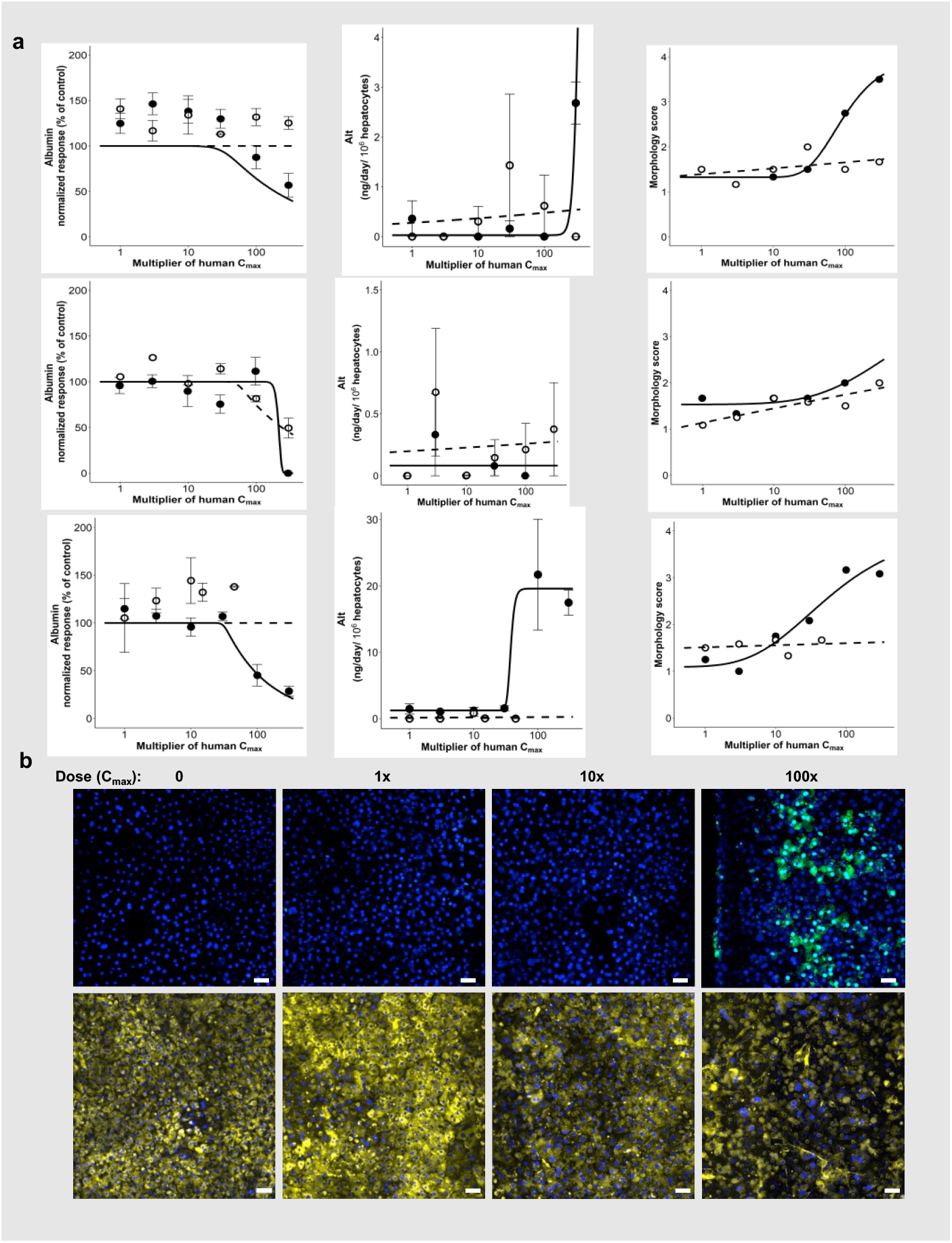
Detection of drug concentration-dependent toxicity and liver injury. **a)** Albumin (left), ALT (middle) and morphological injury score (right) for known non-toxic drugs (open circles) and their closely related toxic partner compounds (open circles) measured on day 3. Clozapine and olanzapine are shown at the top, troglitazone and pioglitazone below, and trovafloxacin and levofloxacin in the bottom row. **b)** Immunofluorescence microscopic images showing concentration-dependent increases in caspase 3/7 staining (green) indicative of apoptosis after treatment with trovafloxacin at 0,1,10, and 100 times the unbound human C_max_ for 7 days (top). The bottom panel shows a concentration-dependent decrease in TMRM staining (yellow) indicative of mitotoxicity in response to treatment with sitaxsentan under similar conditions.

**Table 2.** Data for the matched pair analysis proposed by IQ MPS guidelines. Data from donors one, two and three are presented in terms of multiples of the unbound human C_max_ for each drug, to ease comparison within each pair. Included is the concentration causing a 50% reduction in albumin production, or the lowest concentration causing an increase in ALT protein or cellular morphology score. The dash indicates that the drug was not tested in donor 2 or 3. Apoptosis was defined as a concentration-dependent increase in caspase 3/7 staining, mitotoxicity was defined as a concentration-dependent reduction in TMRM staining and steatosis reflects an increase in Adipored staining. Representative images are contained in Supplementary Figure S3.

| Drug | Albumin IC<sub>50</sub> |  |  | ALT |  |  | Morphology |  |  | IF imaging |  |  |
| --- | --- | --- | --- | --- | --- | --- | --- | --- | --- | --- | --- | --- |
|  | Donor 1 | Donor 2 | Donor 3 | Donor 1 | Donor 2 | Donor 3 | Donor 1 | Donor 2 | Donor 3 | Donor 1 | Donor 2 | Donor 3 |
| **Clozapine** | 33 | 91 | 67 | 300 | > 300 | 100 | 100 | 100 | 100 | Apoptosis | Apoptosis | Apoptosis |
| **Olanzapine** | > 300 | > 300 | > 300 | > 300 | > 300 | > 300 | > 300 | > 300 | > 300 | Apoptosis | No findings | No findings |
| **Fialuridine** | < 0.1 | < 1 | - | > 300 | > 300 | - | > 300 | > 300 | - | No findings | Steatosis | - |
| **FIRU** | > 300 | > 300 | - | > 300 | > 300 | - | > 300 | > 300 | - | No findings | Steatosis | - |
| **Nefazodone** | 140 | - | - | 300 | - | - | 300 | - | - | No findings | - | - |
| **Buspirone** | 282 | - | - | > 300 | - | - | > 300 | - | - | Apoptosis | - | - |
| **Sitaxsentan** | 4.4 | 164 | > 100 | 300 | 300 | 100 | 100 | 100 | 100 | Mitotoxicity | Mitotoxicity | Mitotoxicity |
| **Ambrisentan** | > 300 | > 1000 | > 300 | > 300 | > 1000 | > 300 | > 300 | > 1000 | > 300 | No findings | No findings | No findings |
| **Tolcapone** | 3 | 39 | - | 100 | 30 | - | 10 | 100 | - | Mitotoxicity | Mitotoxicity | - |
| **Entacapone** | > 300 | - | - | > 300 | - | - | > 300 | - | - | No findings | - | - |
| **Troglitazone** | 24 | 122 | - | 300 | 300 | - | 300 | 300 | - | Apoptosis | Apoptosis | - |
| **Pioglitazone** | 277 | > 300 | - | > 300 | > 300 | - | > 300 | > 300 | - | Apoptosis | No findings | - |
| **Trovafloxacin** | 82 | 78 | 31 | 100 | 100 | 30 | 100 | 100 | 30 | Apoptosis | Apoptosis | Apoptosis |
| **Levofloxacin** | > 15 | > 45 | > 45 | > 15 | > 45 | > 45 | > 15 | > 45 | > 45 | No findings | No findings | No findings |

Together, these data support the Liver-Chip’s value as a predictor of drug-induced toxicity in the human liver and demonstrate that this experimental system meets the basic IQ MPS criteria for preclinical model functionality. However, in addition to the seven matched pairs, the IQ MPS guidelines require that an effective human MPS DILI model can predict liver responses to six additional small-molecule drugs associated with clinical DILI. We only analyzed the effects of five of these drugs (diclofenac, asunaprevir, telithromycin, zileuton, and lomitapide) because the reported mechanism of toxicity of one of them (pemoline) is immune-mediated hypersensitivity^35^, and this would potentially require a more complex configuration of the Liver-Chip containing additional immune cells. We also were unable to obtain one of the uggested drugs, mipomersen, from any commercial vendor; however, we tested lomitapide as an alternate, as both produce steatosis by altering triglyceride export, and lomitapide is known to induce elevated ALT levels^36^.

Results obtained with these drugs are presented in Table 3, with toxicity values indicating the lowest concentration at which toxicity was detected. Lomitapide was highly toxic when tested over all included concentrations down to 0.1x human plasma C_max_, with all Liver-Chips showing signs of toxicity following five days of dosing. While telithromycin displayed a decrease in albumin along with a concomitant increase in ALT and morphological injury score, diclofenac and asunaprevir induced concentration and time-dependent changes in albumin and injury scores, but no elevation of ALT was seen with these drugs. Hepatotoxicity was also confirmed with immunofluorescence microscopy, which revealed apoptosis-mediated cell death following exposure to diclofenac, asunaprevir, or telithromycin. However, the Liver-Chip was unable to detect hepatotoxicity caused by zileuton, a treatment intended for asthma. The exact mechanism of toxicity of zileuton is unknown, but it likely involves production of intermediate reactive metabolites due to oxidative metabolism by the cytochrome P450 isoenzymes 1A2, 2C9, and 3A4^37^. Although zileuton is >93% plasma protein bound^38^, we do not believe this was responsible for the lack of toxicological effect, as we were able to detect toxicities induced by other highly protein-bound drugs in the test set.

**Table 3.** Results obtained with the expanded drug list. Data are presented as the lowest unbound human C_max_ multiplier causing a 50% reduction in albumin production, or the lowest concentration causing an increase in ALT release or morphology score. The dash indicates that the drug was not tested in donor 2. Apoptosis was defined as a concentration-dependent increase in caspase 3/7 staining and mitotoxicity was defined as a concentration-dependent reduction in TMRM staining. Representative images are contained in Supplementary Figure S3.

| Drug | Albumin IC<sub>50</sub> |  | ALT |  | Morphology |  | IF imaging |  |
| --- | --- | --- | --- | --- | --- | --- | --- | --- |
|  | Donor 1 | Donor 2 | Donor 1 | Donor 2 | Donor 1 | Donor 2 | Donor 1 | Donor 2 |
| Asunaprevir | 190 | - | > 1000 | - | 1000 | - | Apoptosis | - |
| Benoxaprofen | 8 | 20 | > 100 | 100 | < 0.1 | 100 | Apoptosis | Apoptosis |
| Beta-Estradiol | > 300 | - | > 300 | - | > 300 | - | No findings | - |
| Chlorpheniramine | > 300 | - | > 300 | - | > 300 | - | Apoptosis | - |
| Diclofenac | 384 | - | > 1000 | - | 1000 | - | Apoptosis | - |
| Labetalol | 26 | 40 | > 100 | 100 | > 100 | 100 | Apoptosis | Apoptosis |
| Lomitapide | 4 | - | > 1000 | - | < 0.1 | - | No findings | - |
| Simvastatin | < 0.1 | < 10 | > 300 | > 300 | > 300 | > 300 | No findings | Apoptosis |
| Stavudine | > 60 | 132 | 300 | > 300 | 300 | 300 | No findings | Mitotoxicity |
| Tacrine | 76 | > 25 | > 300 | > 1000 | > 300 | > 1000 | Apoptosis | Apoptosis |
| Telithromycin | 13 | - | 30 | - | 100 | - | Apoptosis | - |
| Ximelagatran | 32 | > 300 | 30 | > 300 | 100 | 300 | No findings | No findings |
| Zileuton | > 100 | 210 | > 100 | > 300 | > 100 | > 300 | No findings | No findings |

### Improved sensitivity for DILI prediction compared to spheroids and animal models

After fulfilling the major criteria of the IQ MPS affiliate guidelines, we considered the Liver-Chip to be qualified as a suitable tool to predict DILI during preclinical drug development. However, we wished to also quantify the performance of the Liver-Chip in the predictive toxicology context. To do so, we expanded the drug test to include eight additional drugs (benoxaprofen, beta-estradiol, chlorpheniramine, labetalol, simvastatin, stavudine, tacrine, and ximelagatran) that were found to induce liver toxicity clinically, despite having gone through standard preclinical toxicology packages involving animal models prior to first-in-human administration. Importantly, the toxicities of these 8 drugs have been shown to be poorly predicted by hepatic spheroids^26,27^.

We proceeded to quantify any observed toxicity across the combined and blinded 27-drug set as a margin of safety (MOS)-like figure by taking the ratio of the minimum toxic concentration observed to the clinical C_max_. We obtained the minimum toxic concentration by taking the lowest concentration identified by each of the primary endpoints—i.e., IC_50_ values for the decrease in albumin production, the lowest concentration at which we observed an increase in ALT protein, and the lowest concentration at which we observed injury via morphology scoring (Table 4). Minimum toxic concentrations generally corresponded to day seven values, although day three values were occasionally lower. We then compared the MOS-like figures against a threshold value of 50 to categorize each compound as toxic or non-toxic, as previously reported for 3D hepatic spheroids, which used a similar threshold^26^. Analyzed in this manner, we found that, in addition to the drugs assessed as part of the IQ MPS-related analysis, the Liver-Chip correctly determined labetalol and benoxaprofen to be hepatotoxic, a response that was consistent across donors one and two and was indicated primarily by a reduction in albumin production. However, we found that simvastatin and ximelagatran were only toxic in one of the donors tested, again showing the importance of including multiple donors during the risk assessment process. Overall, the Liver-Chip correctly predicted toxicity in 12 out of 15 toxic drugs that were tested using two donors, yielding a sensitivity of 80% on this drug set. This was almost double the sensitivity of 3D hepatic spheroids for the same drug set (42%) based on previously published data^26,39,40^, a preclinical model that is currently widely used in pharma and was only able to correctly identify 8 out of the 19toxic drugs in the set (Table 5a). Importantly, the Liver-Chip also did not falsely mark any drugs as toxic (specificity of 100%), whereas the 3D hepatic spheroids did (only 67% specificity)^26^; such false positives can significantly limit the usefulness of a predictive screening technology because of the profound consequences of erroneously failing safe and effective compounds. Interestingly, the three drugs not detected by Liver-Chip—levofloxacin, stavudine, and tacrine—were not detected as toxic drugs in spheroids either, suggesting that the Liver-Chip may subsume the sensitivity of spheroids and that their toxicities could involve other cells or tissues not present in these models. It is important to note that each of the toxic drugs tested was historically evaluated using animal models, and in each case the considerations and thresholds were deemed relevant for that drug to have an acceptable therapeutic window and thus progress into clinical trials. The ability of the Liver-Chip to flag 80% of these drugs for their DILI risk at their clinical concentrations represents a significant improvement in model sensitivity that could drive better decision making in preclinical development.

**Table 4.**
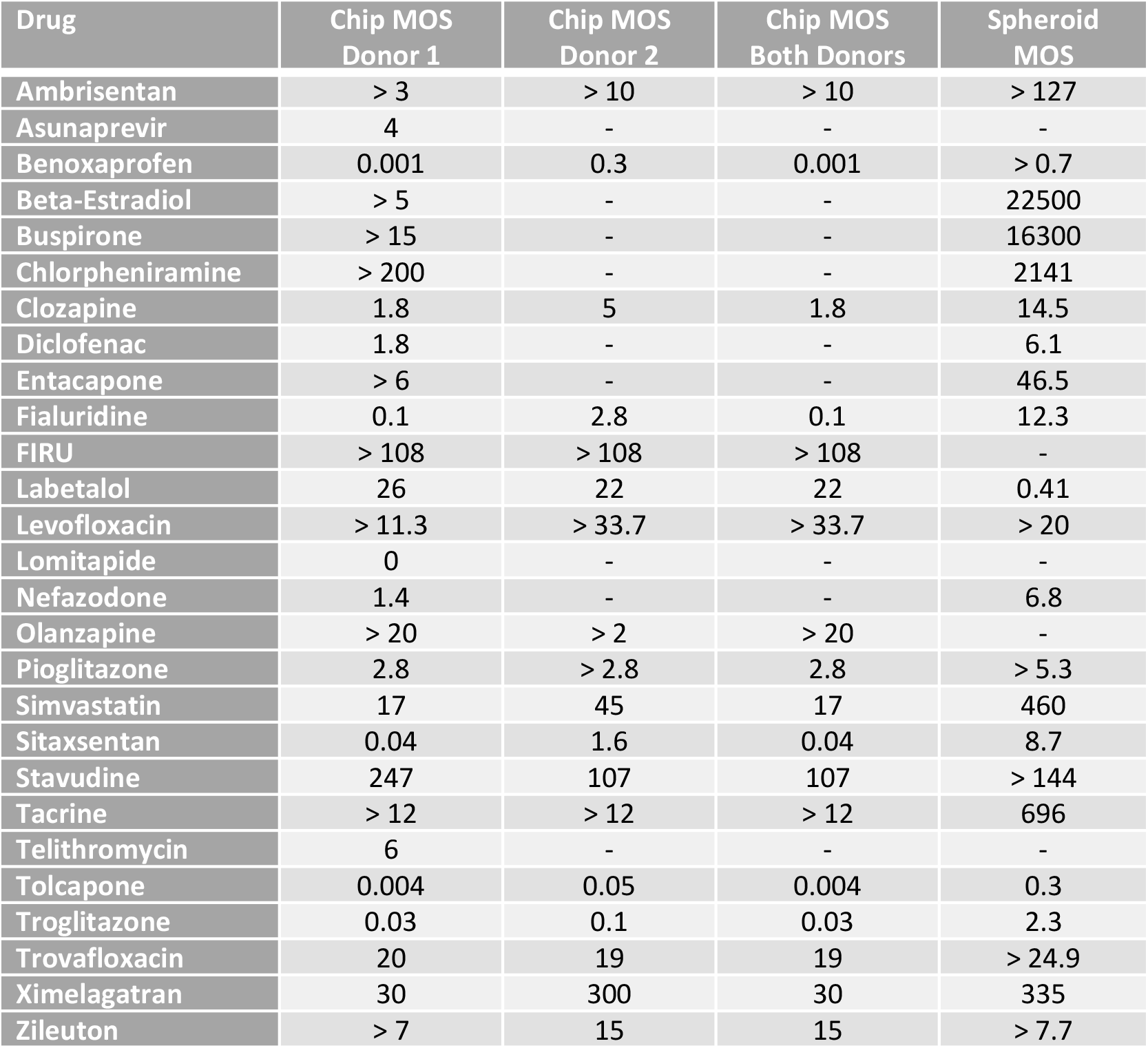
Calculation of Margin of Safety (MOS)-Like Figures. The analysis was carried out as described previously26, that is, the free IC_50_ concentration of the drug tested in the assay divided by the total concentration of drug in human plasma at C_max_.

| Drug | Chip MOS Donor 1 | Chip MOS Donor 2 | Chip MOS Both Donors | Spheroid MOS |
| --- | --- | --- | --- | --- |
| Ambrisentan | > 3 | > 10 | > 10 | > 127 |
| Asunaprevir | 4 | - | - | - |
| Benoxaprofen | 0.001 | 0.3 | 0.001 | > 0.7 |
| Beta-Estradiol | > 5 | - | - | 22500 |
| Buspirone | > 15 | - | - | 16300 |
| Chlorpheniramine | > 200 | - | - | 2141 |
| Clozapine | 1.8 | 5 | 1.8 | 14.5 |
| Diclofenac | 1.8 | - | - | 6.1 |
| Entacapone | > 6 | - | - | 46.5 |
| Fialuridine | 0.1 | 2.8 | 0.1 | 12.3 |
| FIRU | > 108 | > 108 | > 108 | - |
| Labetalol | 26 | 22 | 22 | 0.41 |
| Levofloxacin | > 11.3 | > 33.7 | > 33.7 | > 20 |
| Lomitapide | 0 | - | - | - |
| Nefazodone | 1.4 | - | - | 6.8 |
| Olanzapine | > 20 | > 2 | > 20 | - |
| Pioglitazone | 2.8 | > 2.8 | 2.8 | > 5.3 |
| Simvastatin | 17 | 45 | 17 | 460 |
| Sitaxsentan | 0.04 | 1.6 | 0.04 | 8.7 |
| Stavudine | 247 | 107 | 107 | > 144 |
| Tacrine | > 12 | > 12 | > 12 | 696 |
| Telithromycin | 6 | - | - | - |
| Tolcapone | 0.004 | 0.05 | 0.004 | 0.3 |
| Troglitazone | 0.03 | 0.1 | 0.03 | 2.3 |
| Trovafloxacin | 20 | 19 | 19 | > 24.9 |
| Ximelagatran | 30 | 300 | 30 | 335 |
| Zileuton | > 7 | 15 | 15 | > 7.7 |

We examined each of the toxic drugs that were missed by the Liver-Chips to identify opportunities for future improvement. Using the threshold of 50 for determining toxicity, which we chose to compare our results to those from a past hepatic spheroid study, led to stavudine being classified as a false negative. In fact, Liver-Chips do capture stavudine’s toxicity at a higher threshold without introducing false positives, as described below. Tacrine is a reversible acetylcholinesterase inhibitor that undergoes glutathione conjugation by the phase II metabolizing enzyme glutathione S-transferase in liver. Polymorphisms in this enzyme can impact the amount of oxidative DNA damage, and the M1 and T1 genetic polymorphisms are associated with greater hepatotoxicity^41^. It is not known if either of the two hepatocyte donors used in this investigation have these polymorphisms, but the Liver-Chip was able to detect increased caspase 3/7 staining—indicative of apoptosis at the highest tested concentrations—although these changes were not associated with any release of ALT or decline in albumin. Levofloxacin, a fluoroquinolone antibiotic, was proposed by the IQ MPS affiliate as a lesser hepatotoxin compared to its structural analog trovafloxacin, but it is classified as high clinical DILI concern in Garside’s DILI severity category labeling^42^. Indeed, there are documented reports of hepatotoxicity with levofloxacin, but these occurred in individuals aged 65 years and above,^43^ and a post-market surveillance report documented the incidence of DILI to be less than 1 in a million people^44^. It is therefore reasonable to assume that the negative findings in both the Liver-Chip and spheroids may correctly represent clinical outcome.

### Accuracy improved by accounting for drug-protein binding

When calculating the MOS-like values in the preceding section, we followed the published methods used for evaluating 3D hepatic spheroids^26^, but these do not consider protein binding. Because the fundamental principles of drug action dictate that free (unbound) drug concentrations drive drug effects, we explored an alternative methodology for calculating the MOS-like values by accounting for protein binding using a previously reported approach^31^. Accordingly, we reanalyzed the findings for the 27 drugs in our study by accounting for protein binding. We compared the free fraction of drug concentration dosed in the Liver-Chip employing a medium containing 2% fetal bovine serum to the free fraction of the plasma C_max_. By reanalyzing the Liver-Chip results using this approach and setting the threshold value to 375 (which we selected to maximize sensitivity while avoiding false positives), we obtained improved chip performance: a true positive rate (sensitivity) of 77% and 73% in donors one and two, respectively, and a true negative rate (specificity) of 100% in both donors (Table 5b). Importantly, the sensitivity increased to 87% when including the 18 drugs tested in both donors, and this enabled detection of stavudine’s toxicity. Applying the same analysis to spheroids and similarly selecting a threshold to maximize sensitivity while maintaining 100% specificity yielded a sensitivity of only 47%. Remarkably, the Spearman correlation between the two-donor Liver-Chip assay and the Garside DILI severity scale yielded a value of 0.78 when using the protein binding-corrected analysis, whereas it was only 0.43 when using the lower threshold. Thus, the protein-binding-corrected approach not only produces higher sensitivity but also rank-orders the relative toxicity of drugs in a manner that corresponds better with the DILI severity observed in the clinic. This observation supports the validity of this analysis approach and its superiority over the uncorrected version. In short, these results provide further confidence that the Liver-Chip is a highly predictive DILI model and is superior in this capacity to other currently used approaches.

### The economic value of more predictive toxicity models in preclinical decision making

In addition to increasing patient safety, better prediction of candidate drug toxicity can improve the economics of drug development by reducing clinical trial attrition and increasing pharma research and development (R&D) productivity. We sought to quantify the potential economic impact of the Liver-Chip resulting from its enhanced predictive validity by constructing an economic value model of drug development that captures decision quality during preclinical development (Figure 4). We describe the structure of this model in the Methods section and provide an interactive form of the full model in the Supplementary Materials.

**Figure 4.**
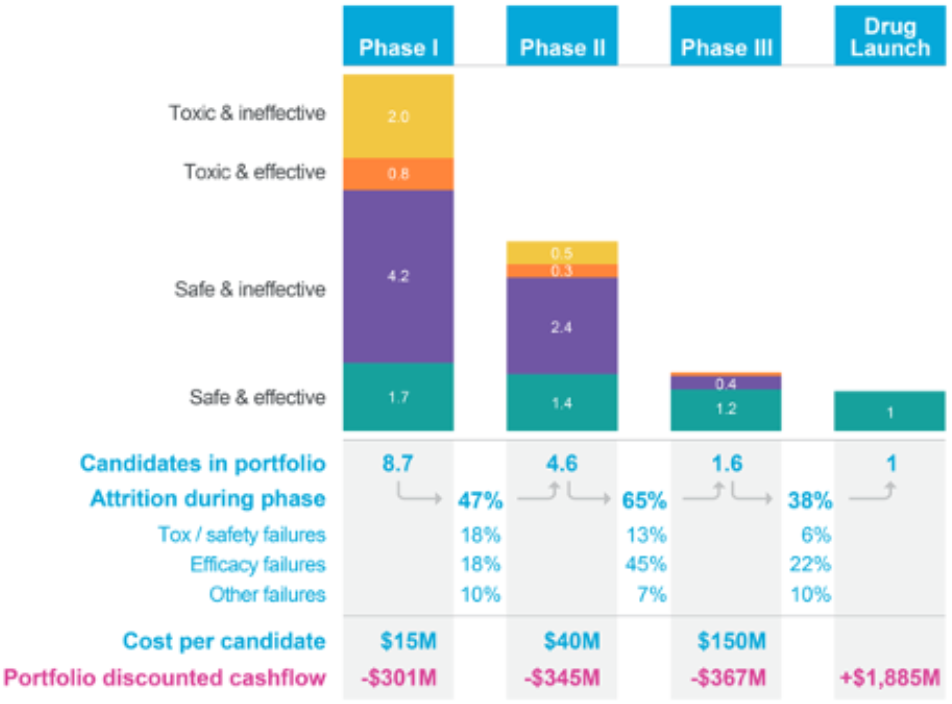
Economic value model for assessing the financial impact of improved preclinical testing. Illustrated is the model’s “base case”, which tracks a representative portfolio of candidate drugs as it progresses and erodes through clinical trials, culminating in a single drug approval. The model bases phase-by-phase attrition rates (“attrition during phase”), discovery and preclinical costs, development costs (“cost per candidate”) and cost of capital on Paul et al. (2010)^5^ to compute a portfolio-wide discounted cashflow. In contrast with prior approaches, the model tracks the underlying causes of clinical trial failure (safety-related, efficacy-related, and other failures) using parameters derived from literature^7,9, 46,72^, a feature that permits us to determine the composition of the drug portfolio in each stage of development in terms of candidates that are safe and effective, safe and ineffective, unsafe and effective, and unsafe and ineffective, as illustrated. Improvements in the predictive validity of preclinical safety testing can be captured through their impact on the makeup of the portfolio entering Phase I clinical trials: better preclinical safety testing reduces the proportion of unsafe drugs that enter the clinic relative to the “base case”; the model permits analyzing the impact of such changes on the discounted cashflow and the portfolio’s profitability. The model is provided in full in the supplementary materials as a formula-driven Microsoft Excel file.

To estimate the economic impact of incorporating the Liver-Chip into preclinical research, we observed that DILI currently accounts for 13% of clinical trial failures that are due to safety concerns^46^. The present study revealed that the Liver-Chip, when used with two donors and analyzed with consideration for protein binding, provides a sensitivity of 87% when applied to compounds that evaded traditional safety workflows. Combining these figures suggests that using the human Liver-Chip to test for DILI risk could lead to 11.3% fewer toxic drugs entering clinical trials. We modeled this improvement by correspondingly lowering the model’s false negative rate (FNR) parameter that describes the toxicology testing that occurs between preclinical testing and Phase I clinical trials. We then computed the net present value (NPV) of the new simulated portfolio and compared it to the NPV of the base case to capture the increase in R&D productivity as described in the Methods section. This computation resulted in a predicted NPV uplift of 2.8% (1.9% - 3.1%, CI 95%) due to the incorporation of the Liver-Chip in DILI prediction (Supplementary Table S2 lists results for a broad range of FNR values).

We next estimated the impact of this uplift on the broader small-molecule drug-development industry by applying it the global Pharma investment in R&D, which currently approximates $196m per year^73^ of which around 56%^74^ is related to small-molecule drugs. Remarkably, the model predicts that utilizing the Liver-Chip across all small-molecule drug development programs for this single context of use could generate the industry around $3 billion annually due to increased R&D productivity ($2.1B - $3.4B, CI 95%). Since the model relies on historical attrition rates and costs, we assessed the robustness of the above predictions with respect to the model’s inputs by performing a sensitivity analysis as described in the Methods section. This analysis revealed that model outputs vary in a near-linear fashion across reasonable input parameter sets, thereby retaining the qualitative conclusions regardless of parameter choices. The details of this analysis and its results are included in the model as part of the Supplementary Materials.

The economic model also permits us to estimate the financial impact of Organ-Chip technology as the predictive validity of additional toxicology models is evaluated similarly to our work here on the Liver-Chip. We were particularly interested in the potential impact of four additional Organ-Chips that address the remaining top causes of safety failures— cardiovascular, neurological, immunological, dermatological, and gastrointestinal toxicities, which together with DILI account for 80% of trial failures due to safety concerns^46^. If we assume similar sensitivity for these four additional models as we found for the Liver-Chip (87%), the model estimates that Organ-Chip technology could generate the industry over $24 billion annually through increased R&D productivity. These figures present a compelling economic incentive for the adoption of Organ-Chip technology alongside considerations of patient safety and the ethical concerns of animal testing.

## Discussion

Organ-Chip technology has tremendous potential to revolutionize drug discovery and development^47^, and many major pharmaceutical companies have already invested in the technology, but routine utilization is limited^48^. This may be due to several factors, but the overriding fact is either that there has not been an end-to-end investigation showing that Organ-Chips replicate human biological responses in a robust and repeatable manner, that its performance exceeds that of existing preclinical models, or that there is a way to implement the technology into routine drug screening projects. This investigation directly addressed these three concerns. Furthermore, the broader stakeholder group—especially budget holders—need assurance that there will be a return on investment and that such technologies will help reverse the pharmaceutical industry’s widely documented productivity crisis.

To counteract the R&D productivity crisis^49^, the pharmaceutical industry is seeking physiologically relevant models that can be incorporated into the early-stage drug discovery programs in which the cost of attrition is lower and, ultimately, the quality of drug candidates progressing into the clinic will be higher^50,51^. Organ-on-a-chip technology utilizes microengineering to develop physiologically complex, human-relevant models; therefore, this technology should be implemented into programs to achieve this goal. To date, there has been no systematic evaluation of the validity of Organ-Chips or any other MPS for DILI prediction against criteria designed by a third party of experts. To our knowledge, no MPS has been evaluated against 27 small-molecule drugs in a single study involving three different human donors and hundreds of experiments, making this study the largest reported evaluation of Organ-Chip performance. The Liver-Chip has demonstrated that it can correctly distinguish toxic drugs from their non-toxic structural analogs, and, across a blinded set of 27 small molecules, has a true positive rate of 87%, a specificity of 100%, and a Spearman correlation of 0.78 against the Garside DILI severity scale when two donors are used, and data are corrected for protein binding. Importantly, these data were independently verified by two external toxicologists. Said differently, the Liver-Chip detected nearly 7 out of every 8 drugs that proved hepatoxic in clinical use despite having been deemed to have an appropriate therapeutic window by animal models; the Liver-Chip similarly detected 2 out of 4 such drugs that were additionally missed by 3D hepatic spheroids. Hence, we believe that these findings advocate the routine use of the human Liver-Chip in drug discovery programs to enhance the probability of clinical success while improving patient safety (Figure 5). This would be achieved by more-accurately categorizing risk associated with a candidate drug to provide valuable data to support a ‘weight-of-evidence’ argument both for entry into the clinic as well as for starting dose in phase I. Such added evidence could potentially remove any safety factor applied because of a liver finding in an animal model^52,53^. In turn, this would reduce overall cost and time in the preclinical development process.

**Figure 5.**
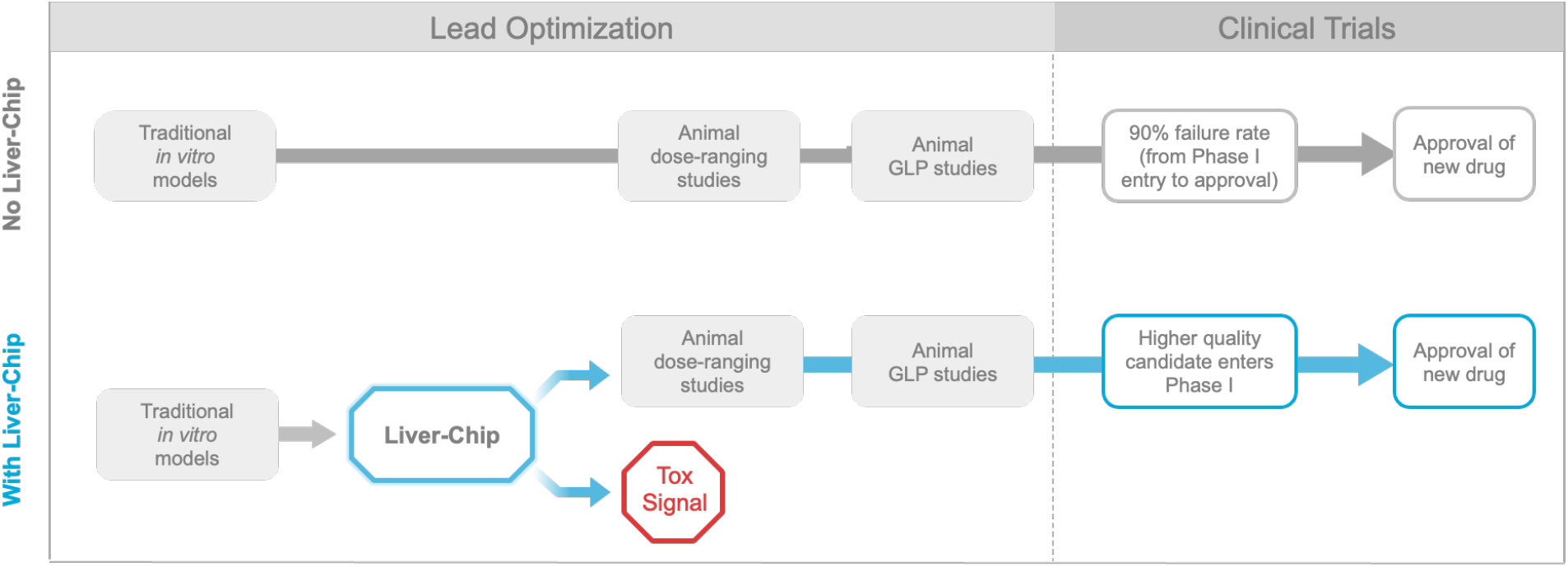
Proposed positioning of the Liver-Chip within a typical pharma pre-clinical workflow. Typically, pharma utilizes a series of *in vitro* tests to guide chemical optimization ahead of animal testing. Promising drug candidates then progress to dose-range finding studies ahead of the required studies to enable regulatory approval to enter clinical trial. With the data presented in this investigation, Liver-Chip would be best placed in between the *in vitro* tests and dose-range finding animal studies. A drug candidate that did not show toxicity in the Liver-Chip, would increase confidence of the scientist that it can pass through animal testing without a liver toxicity flag and proceed into the clinic with a lower likelihood of clinical hepatic signals. A drug candidate that did show toxicity in the Liver-Chip would encourage scientists to stop and think about the relevance of the toxicity to the therapeutic indication and whether there was a potential margin between this finding and the exposure required for clinical efficacy. This would continue to increase the confidence that candidate drugs are entering the phase I clinical trial process with a greater likelihood of approval and may also reduce animal usage by not conducting dose-range finding or regulatory studies.

A unique feature of this work is the demonstration of the throughput capability of Organ-Chip technology using automated culture instruments, as a total of 870 chips were created and analyzed. In terms of establishing effective workflows, scientists were placed into three teams: the first team prepared the drug solutions and supplied them in a blinded manner to the second team. The second team seeded, maintained, and dosed the Liver-Chips while carrying out various morphological, biochemical, and genetic analyses at the end of the experiment. The third team collected the effluents and performed real-time analyses of albumin and ALT as well as terminal immunofluorescence imaging using an automated confocal microscope (Opera Phenix; Perkin Elmer). In this manner, we were able to analyze and report the hepatotoxic effects of 27 drugs in 870 Liver-Chips that used cells from three human donors in a period of 20 weeks.

Based on this experience, we believe that the Liver-Chip could be employed in the drug development pipeline during the lead optimization phase where projects have identified three-to-five chemical compounds that have the potential to become the candidate drug. If data emerge showing that a chemical compound produces a toxic signal in the Liver-Chip, this will indicate to toxicologists that there is a high (∼87%) probability that the compound would similarly cause toxicity in humans. This, in turn, would enable scientists to deprioritize these compounds from early *in vivo* toxicology studies (such as the maximum tolerated dose/dose range finding study) and, consequently, reduce animal usage and advance the “fail early, fail fast” strategy. Importantly, the absence of false positives strengthens the argument that the Liver-Chips should also be adopted within the early discovery phase, as stopping drug candidates that are falsely determined to be toxic by less-robust preclinical models could result in good therapeutics never reaching patients.

Despite these positive findings, it should be acknowledged that the current chip material (PDMS) used in the construction of the Liver-Chip may be problematic for a subset of small molecules that are prone to non-specific binding. Although this study demonstrates that the material binding issue does not in practice significantly reduce the predictive value of the Liver-Chip DILI model, work is currently underway to develop chips using materials that have a lower binding potential. Until such a chip is available, we recommend users assess potential PDMS binding using an acellular chip and measuring drug in the effluent channel using LC/MS to enable adjustment of workflow if required. It should also be recognized that many pharmaceutical companies have diversified portfolios, with only 40 to 50% now being small molecules. Consequently, further investigation of the Liver-Chip performance against large molecules and biologic therapies should be carried out. Integration of resident and circulating immune cells should add even greater predictive capability.

Finally, predictive models that demonstrate concordance with clinical outcomes should provide scientists and corporate leadership with greater confidence in decision-making at major investment milestones. Impressively, our economic analysis revealed $3 billion in improved R&D productivity that could be generated by replacing or supplementing existing preclinical *in vitro* models with human Liver-Chips for this single context of use (DILI prediction). Moreover, an additional productivity value of $24 billion could be gained if a similar approach is used to develop predictive models for the other most common toxicities that result in drug attrition. Taken together, these results suggest that Organ-Chip technology has great potential to benefit drug development, improve patient safety, and enhance pharmaceutical industry productivity and capital efficiency. This work also provides a starting point for other groups that hope to validate their MPS models for integration into commercial drug pipelines.

## Methods

### Cell Culture

Cryopreserved primary human hepatocytes, purchased from Gibco (Thermo Fisher Scientific), and cryopreserved primary human liver sinusoidal endothelial cells (LSECs), purchased from Cell Systems, were cultured according to their respective vendor/Emulate protocols. The LSECs were expanded at a 1:1 ratio in 10-15 T-75 flasks (Corning) that were pre-treated with 5mL of Attachment Factor (Cell Systems). Complete LSEC medium includes Cell Systems medium with final concentrations of 1% Pen/Strep (Sigma), 2% Culture-Boost (Cell Systems), and 10% Fetal Bovine Serum (FBS) (Sigma). Media was refreshed daily until cells were ready for use. Cryopreserved human Kupffer cells (Samsara Sciences) and human Stellate cells (IXCells) were thawed according to their respective vendor/Emulate protocols on the day of seeding. See Supplementary Table S1 for further information.

### Liver Chip Microfabrication and Zoë® Culture Module

Each chip is made from flexible polydimethylsiloxane (PDMS), a transparent viscoelastic polymer material. The chip compartmental chambers consist of two parallel microchannels that are separated by a porous membrane containing pores of 7 µm diameter spaced 40 µm apart.

On Day -6, chips were functionalized using Emulate proprietary reagents, ER-1 (Emulate reagent: 10461) and ER-2 (Emulate reagent: 10462), mixed at a concentration of 1 mg/mL prior to application to the microfluidic channels of the chip. The platform is then irradiated with high power UV light (peak wavelength: 365nm, intensity: 100 µJ/cm^2^) for 20 minutes using a UV oven (CL-1000 Ultraviolet Crosslinker AnalytiK-Jena: 95-0228-01). Chips were then coated with 100 µg/mL Collagen I (Corning) and 25 µg/mL Fibronectin (ThermoFisher) in both channels. The top channel was seeded with primary human hepatocytes on Day -5 at a density of 3.5 × 10^6^ cells/mL. Complete hepatocyte seeding medium contains Williams’ Medium E (Sigma) with final concentrations of 1% Pen/Strep (Sigma), 1% L-GlutaMAX (Gibco), 1% Insulin-Transferring-Selenium (Gibco), 0.05 mg/mL Ascorbic Acid (Sigma), 1 µM dexamethasone (Sigma), and 5% FBS (Sigma). After four hours of attachment, the chips were washed by gravitational force. Gravity wash consisted of gently pipetting 200 µL fresh medium at the top inlet, allowing it to flow through, washing out any unbound cells from the surface, and inserting a pipette tip on the outlet of the channel.

On Day -4, a hepatocyte Matrigel overlay procedure was executed with the purpose of promoting a three-dimensional matrix for the hepatocytes to grow in an ECM sandwich culture. The hepatocyte overlay and maintenance medium contains Williams’ Medium E (Sigma) with final concentrations of 1% Pen/Strep (Sigma), 1% L-GlutaMAX (Gibco), 1% Insulin-Transferrin-Selenium (Gibco), 50 µg/mL Ascorbic Acid (Sigma), and 100 nM Dexamethasone (Sigma). On Day -3, the bottom channel was seeded with LSECs, stellate cells and Kupffer cells, further known as non-parenchymal cells (NPCs). NPC seeding medium contains Williams’ Medium E (Sigma) with final concentrations of 1% Pen/Strep (Sigma), 1% L-GlutaMAX (Gibco), 1% Insulin-Transferrin-Selenium (Gibco), 50 µg/mL Ascorbic Acid (Sigma), and 10% FBS (Sigma). LSECs were detached from flasks using Trypsin (Sigma) and collected accordingly. These cells were seeded in a mixture volume ratio of 1:1:1 with LSECs at a density range of 9-12 × 10^6^ cells/mL, stellates at a density of 0.3 × 10^6^ cells/mL, and Kupffer cells at 6 × 10^6^ cells/mL followed by a gravity wash four hours post-seeding.

On Day -2, chips were visually inspected under the ECHO microscope (Discover Echo, Inc.) for cellular maturation and attachment, healthy morphology, and a tight monolayer. The chips that passed visual inspection had both channels washed with their respective media, leaving a droplet on top. NPC maintenance media was composed of the same components prior, with a reduction of FBS to 2%. To minimize bubbles within the system, one liter of complete, warmed top and bottom media was added to Steriflip-connected tubes (Millipore) in the biosafety cabinet. All media was then degassed using a -70 kPa vacuum (Welch) and stored in the incubator until use. Pods were primed twice with 3 mL of degassed media in both inlets, and 200 µL in both outlets. Chips were then connected to pods via liquid-to-liquid connection. Chips and pods were placed in the Zoë^®^ (Emulate Inc.) for their first regulate cycle, which minimizes bubbles within the fluidic system by increasing the pressure for two hours. After this, normal flow resumed at 30 µL/h. On Day -1, Zoës^®^ were set to regulate once more.

### Experimental Setup

The 870-chip experiment was carried out in five consecutive cycles (herein referred to as Cycles 1 through 5) to test a selection of 27 drugs at varying concentrations relative to the average therapeutic human C_max_ obtained from literature (Supplementary Table S2). Cycles 1 to 4 tested 6-8 concentrations in duplicate for each of 10-13 drugs. To determine which sampling strategy was optimal for cycles 1 to 4 (16 doses x 1 replicate, 8×2, 5×3, or 4×4), we generated 3 different “dose-response” synthetic datasets, each distorted by different noise levels (low, medium, or high). For each of these datasets, we performed curve-fitting analysis and calculated the Root Mean Square Error between the “true” and estimated IC_50_ parameter. The analysis results showed that, for all noise levels, the 16×1 sampling strategy marginally outperformed the 8×2. However, to ensure at least two replicates per concentration, the 8×2 strategy was selected. Some drugs were also repeated across cycles (on different donors or at different concentrations) to ensure experimental robustness. However, to replicate a more typical study design likely to be carried out by scientists in the pharmaceutical industry, we created cycle 5 with 6 drugs, where each was tested in 4 concentrations in triplicate. For each cycle, chips were dosed with drug over 8 days (referred to as Day 0 through Day 7). Drug preparation, dosing, and analysis teams were divided, creating a double-blind study such that those administering the drugs or performing analyses did not know the name or concentrations of the drugs tested.

### Drug Preparation

The drug dosing concentrations were determined from the unbound human C_max_ of each drug. First, the expected fraction of drug unbound in media with 2% FBS was extrapolated from plasma binding data for each drug. Dosing concentrations were then back calculated such that the unbound fraction in media would reflect relevant multiples of unbound human C_max_ (Supplementary Table S2). For each cycle, concentrations ranged from 0.1 to 1000 times C_max_.

Stock solutions were prepared at 1000 times the final dosing concentration. Drugs in powder form were either weighed out with 1 mg precision or dissolved directly in vendor-provided vials. Sterile DMSO (Sigma) was added to dissolve the drug. The solution was triturated before transferring to an amber vial (Qorpak), which was vortexed (Fisher Scientific) on high for 60 seconds to ensure complete dissolution. A serial dilution was then performed in DMSO to prepare each subsequent 1000X concentration. These stock solutions were then aliquoted in 1.5 mL tubes (Eppendorf) and stored at -20°C until dosing day, allowing a maximum of one freeze-thaw cycle prior to dosing.

All media was made the day prior to chip dosing and stored overnight at 37°C. On the day of chip dosing, one stock aliquot per drug concentration was thawed in a 37°C bead bath. The stocks were then vortexed and inspected to ensure absence of drug particulate. Dosing solutions were prepared by diluting drug stock 1:1000 in top or bottom media to achieve 0.1% DMSO concentration. The dosing solutions were then vortexed and stored at 37°C until dosing.

On the first dosing day (Day 0), all chips were imaged using the ECHO microscope. 500 µL of effluent was collected from all four reservoirs of the pod and placed in a labelled 96-well plate. After collection, all the remaining media was carefully aspirated before dosing with 3.8 mL of corresponding dosing solution. Dosing occurred on study days 0, 2, and 4 for chips flowing at 30 µL/h, and on days 0, 1, 2, 3, 4, 5, and 6 for chips flowing at 150 µL/h. Effluent collection occurred on days 1, 3, and 7.

### Biochemical Assays

Top channel outlet effluents were analyzed to quantify albumin and alanine transaminase (ALT) levels on days 1, 3, and 7, using sandwich ELISA kits (Abcam, Albumin ab179887, ALT ab234578), according to vendor-provided protocols. Frozen (−80°C) effluent samples were thawed overnight at 4°C prior to assay. The Hamilton Vantage liquid handling platform was used to manage effluent dilutions (1:500 for albumin, neat for ALT), preparation of standard curves, and addition of antibody cocktail. Absorbance at 450nm was measured using the SynergyNeo Microplate Reader (BioTek).

As part of cycles 3, 4 and 5, top channel outlet samples from vehicle chips on days 1, 3 and 7 post-drug or vehicle administration were analyzed to quantify urea levels with a urea assay kit (Sigma-Aldrich, MAK006) according to vendor-provided protocol. Frozen (−80°C) effluent samples were thawed overnight at 4°C prior to assay. All samples were diluted 1:5 in assay buffer and mixed with the kit’s Reaction Mix. Absorbance at 570nm was measured using the same automated plate reader.

Effluent samples from vehicle chips and those treated with either trovafloxacin or levofloxacin were thawed overnight at 4°C, and effluents from both channels were analyzed for IL-6 and TNF-alpha levels using MSD U-PLEX kits (Meso Scale Diagnostics, K15067L-2) according to vendor-provided protocols. Samples were added to plates manually at a 1:2 dilution. Plates were read for cytokine release on the MESO QuickPlex SQ 120 (Meso Scale Discoveries).

### Morphological Analysis

At least four to six brightfield images were acquired per chip for morphology analysis. Brightfield images were acquired on the ECHO microscope using these settings: 170% zoomed phase contrast, 50% LED, 38% brightness, 41% contrast, 50% color balance, color on, and 10X objective. Brightfield images were acquired across three fields of view on days 1, 3, and 7 for each cycle. Cytotoxicity classification was performed while acquiring images for both NPCs and hepatocytes. Images were then scored zero to four by blinded individuals (n=2) based on severity of agglomeration of cell debris for both channels. The scoring matrix and representative images have been included in the supplement (Supplementary Figure S2).

At the end of the experiment, cells in the Liver-Chip were fixed using 4% paraformaldehyde (PFA) solution (Electron Microscopy Sciences). Chips were detached from pods and washed once with PBS. The PFA solution was pipetted into both channels and incubated for 20 minutes at room temperature. Afterwards, chips were washed with PBS and stored at 4°C until staining. Following fixation, chips corresponding to low, medium, and high concentrations from each group were cut in half with a razor blade perpendicular to the co-culture channels. One half was used in the following staining protocol, while the other half was stored for future staining. All stains and washes utilized the bubble method, in which a small amount of air is flowed through the channel prior to bulk wash media to prevent a liquid-liquid dilution of the staining solution. The top channel was perfused with 100 µL of AdipoRed (Lonza, PT-7009) diluted 1:40 v/v in PBS labeling lipid accumulation and 100 µL of NucBlue (ThermoFisher, R37605) (100 drops in 50 mL of PBS) to visualize cell nuclei. Following 15 minutes of incubation at room temperature, each channel was washed with 200 µL of PBS (alternating channels, 2x for top and 3x for bottom). As an alternative lipid accumulation marker, 100 µL of HCS LipidTOX Deep Red (ThermoFisher, H34477) was diluted 1:1000 v/v in PBS with NucBlue counterstain and added to the top channel. After a 30-minute incubation at room temperature, the chips were washed with 200 µL of PBS as described previously. Chips were then imaged using the Opera Phenix.

Following lipid and DAPI staining and imaging, chips were stained with a multi-compound resistant protein 2 (MRP2) antibody to visualize the bile canalicular structures characteristic of healthy Liver-Chips. First, chips were permeabilized in 0.125% Triton-X and 2% Normal Donkey Serum (NDS) diluted in PBS (100 µL of solution per channel) and incubated at room temperature for 10 minutes. Then, each channel was washed with 200 µL of PBS (alternating channels, 2x for top and 3x for bottom). Chips were then blocked in 2% Bovine Serum Albumin (BSA) and 10% NDS in PBS (100 µL of solution per channel) and incubated at room temperature for 1 hour. Next, primary antibody Mouse anti-MRP2 (Abcam, ab3373) was prepared 1:100 in the original blocking buffer, diluted 1:4 in PBS. 100 µL of solution was added to each channel, and chips were stored overnight at 4°C. The following day, each channel was washed with 200 µL of PBS (alternating channels, 2x for top and 3x for bottom). Secondary antibody Donkey anti-Mouse 647 (Abcam, ab150107) was prepared 1:500 in original blocking buffer, diluted 1:4 in PBS. 100 µL of solution was added to each channel and chips incubated at room temperature, protected from light, for two hours. Then each channel was washed with 200µL of PBS (alternating channels, 2x for top and 3x for bottom). Chips were imaged immediately or stored at 4°C until ready for imaging on the Opera Phenix.

### Live Staining

Chip replicates designated for live cell imaging were washed with PBS utilizing the bubble method. Chips were then cut in half perpendicular to the co-culture channels. The top chip halves were stained with NucBlue (ThermoFisher, R37605) to visualize cell nuclei and Cell Event Green (ThermoFisher, C10423) to visualize activated caspase 3/7 for apoptosis. This staining panel was prepared in serum-free media (CSC), with NucBlue at 2 drops per mL and Cell Event Green at a 1:500 ratio and perfused through both channels. The bottom chips halves were stained with NucBlue (Thermo) to visualize nuclei and Tetramethylrhodamine, methyl ester (TMRM) (ThermoFisher, I34361) to visualize active mitochondria. This staining panel was prepared in PBS with 5% FBS, with NucBlue at 2 drops per mL and TMRM at a 1:1000 ratio in original blocking buffer, diluted 1:4 in PBS. Chips were incubated in the dark at 37°C for 30 minutes, and then each channel was washed with 200µL of PBS (alternating channels, 2x for top and 3x for bottom). The chips were kept at 37°C, protected from light, until ready for imaging with the Opera Phenix.

### Image Acquisition

Fluorescent confocal image acquisition was performed using the Opera Phenix High-Content Screening System and Harmony 4.9 Imaging and Analysis Software (PerkinElmer). Before acquisition, the Phenix internal environment was set to 37°C and 5% CO_2_. Chips designated for imaging were removed from their plates, wiped on the bottom surface to remove moisture, and placed into the Phenix 12-chip imaging adapter. Whole chips were placed directly into each slot, while top and bottom half chips were matched and combined in one chip slot. Chips were aligned flush with the adapter and one another. Any bubbles identified from visual inspection were washed out with PBS. Once ready, the stained chips were covered with transparent plate film to seal channel ports and loaded into the Phenix. For live imaging, the DAPI (Time: 200ms, Power: 100%), Alexa 488 (Time: 100ms, Power: 100%), and TRITC (Time: 100ms, Power: 100%) lasers were used. For fixed imaging, the DAPI (Time: 200ms, Power: 100%), TRITC (Time: 100ms, Power: 100%), and Alexa 647 (Time: 300ms, Power: 80%) lasers were used. Z-stacks were generated with 3.6µm between slices for 28-32 planes, so that the epithelium was located around the center of the stack. Six fields of view (FOVs) per chip were acquired, with a 5% overlap between adjacent FOVs to generate a global overlay view.

### Image Analysis

Raw images from fixed and live imaging were exported In TIFF format from the Harmony software. Using scripts written for FIJI (ImageJ), TIFFs across 3 color channels and multiple z-stacks were compiled into composite images for each field of view in each chip. The epithelial signal was identified and isolated from the endothelial and membrane signals, and the composite TIFFs were split accordingly. The ideal threshold intensity for each channel in the epithelial “substack” was identified to maximize signal, and the TIFFs were exported as JPEGs for further analysis.

### Gene Expression Analysis

RNA was extracted from chips using TRI Reagent (Sigma-Aldrich) according to the manufacturer’s guidelines. The collected samples were submitted to GENEWIZ (South Plainfield, NJ) for next-generation sequencing. After quality control and RNA-seq library preparation, the samples were sequences with Illumina HiSeq 2×150 system using sequencing depth ∼50M paired-ends reads/sample. Using Trimmomatic v0.36, the sequence reads were trimmed to filter out all poor-quality nucleotides and possible adapter sequences. The remaining trimmed reads were mapped to the Homo sapiens reference genome GRCh38 using the STAR aligner v2.5.2b. Next, using the generated BAM files for each sample, the unique gene hit counts were calculated from the Subread package v1.5.2. It is worth noting that only unique reads within the exon region were counted.

### Statistical Analysis

All statistical analyses were conducted in R^69^ (version 4.1.2) and figures were produced using the R package ggplot2^70^ (version 3.3.5). The dose-response analysis (Figure 3a) was carried out using the popular drc R package developed by Ritz et al.^71^ using the generalized log-logistic dose response model. The error bars in Figures 2d, 2e, 2f and 3a correspond to the standard errors of the mean. The circles in Figure 2d and 2e correspond to the samples used to calculate the corresponding statistics. The analysis of significance in Figures 2d and 2e was performed using unpaired t-test. In Figure 2f the number of samples used were N=3 for donor two, and N=4 for donor three. For both donors, the number of the freshly thawed hepatocyte samples used to estimate the corresponding log2(Fold Change) were N=4. Finally, the analysis of significance in Figures 2f was performed using paired t-test.

### Economic Modeling Approach

An economic model was built to assess the impact of improvements in the predictive validity of preclinical toxicology models on the economics of drug development. This model is provided in full in the supplementary materials as a formula-driven Microsoft Excel file. The model was built by extending the pipeline model of Paul et al. (2010)^5^, which tracks the economics of a representative portfolio of candidate drugs as it progresses and erodes through clinical trials. However, in contrast with conventional models, we followed Scannell & Bosley’s (2016)^4^ approach by modelling attrition as a function of decision quality and candidate quality at each development stage. We modelled safety-related failures, efficacy-related failures, and other failures (e.g., commercial and strategy related) with parameters derived from the literature (primarily from Harrison 2016, ^7,9, 46^). The model comprises a “base case”, which describes an archetypical drug development portfolio that leads to a single drug approval. Development costs, timing, cost of capital and attrition rates were set in line with Paul et al. (2010)^5^. The base case and its parameters are summarized in Figure 4.

An innovative feature of the economic model is that it permits us to determine the makeup of the drug portfolio in each stage of development in terms of candidates that are safe and effective, safe and ineffective, unsafe and effective, and unsafe and ineffective. Additionally, the model’s structure also allows one to estimate decision quality parameters at each stage of the process, such as the false negative rate (FNR) of the toxicity determination – the proportion of toxic drugs erroneously deemed safe. This, in turn, allows one to estimate the financial impact of changes in predictive validity, something that cannot be done directly with conventional attrition-driven pipeline models.

Improvements in the predictive validity of preclinical safety testing can be captured through their effects on the makeup of the portfolio entering Phase I clinical trials: better preclinical safety testing reduces the proportion of unsafe drugs that enter the clinic. Such improvements are captured by reducing the FNR for the toxicology testing that occurs between preclinical development and Phase I trials (we also add Organ-Chip costs to capture the price of added testing). If we keep all other model parameters unchanged, the model captures a cost-avoidance strategy: an approach wherein the ability to predict certain clinical trial failures in advance allows one to start fewer clinical trials (skipping those trials that are bound to fail) to bring one drug to market, as in the base case. However, the ability to have more predictable clinical outcomes is not likely to reduce investment but rather to increase it. This increase in R&D productivity should therefore result at least in maintenance of the investment in clinical testing (if not in its increase), which we conservatively model by setting number of projects entering Phase I to its base case value.

To derive the economic implications of this scenario analysis, we calculate the portfolio’s new net present value (NPV) and evaluate its percentage increase (uplift) over the base case. This NPV uplift represents the increase in R&D productivity caused by the improved testing. The model proceeds to apply this uplift to the world-wide R&D spending on small molecule drug development to estimate the annual financial impact that the increase in R&D productivity may generate.

Because the model is parameterized using historical estimates of attrition rates and their causes, we sought to understand the model’s sensitivity to the exact parameter values. To do this, we performed a mathematical sensitivity analysis for the major input parameters; this analysis is included within the Excel file in the supplementary materials. The analysis demonstrated that reasonable changes in parameter choices retain the model’s qualitative conclusions. This is in part because the model’s output is a percentage uplift relative to the base case, making the model robust in the face of uncertainty in the financials of the base case.

## Acknowledgements

The authors would like to recognize the wider Emulate team for their valuable discussion and support and are grateful for Aaron VanDevender’s critical evaluation and review of the manuscript.

## Author contributions

L.E., D.E.I., D.L., D.V.M., J.D.S. designed the study; A.A., S.A.B., J.T.C., C.V.C., A.R.H., J.J., S.J., K.K.M., M.E.Q., M.K., A.C.R., W.T., M.W., J.V., contributed to drug preparation, execution of the study, downstream processing of images and data collection; J.J., S.J., S.R.J., G.K., V.J.K., C.Y.L., C. L., J.S.R., D.R.T., M.W.,J.F.K.S., chip seeding, maintenance, drug administration, and, chip sampling; C.V.C., S.J., G.K., V.J.K., provided leadership for experimental cycles; L.E., C.L., D.V.M., S.J., M.M. K., J.D.S. contributed to the data analysis; A.R.H., L.E., D.E.I., C.Y.L., K.K.M., M.W. contributed to writing the manuscript; D.L., J.W.S. built the economic model and interpreted the Liver-Chip outcome on R&D productivity; K-J. J., O.I., P.K.M. performed a blinded data review and critically reviewed the manuscript.

## Supplementary Material

**Supplementary Figure S1.**
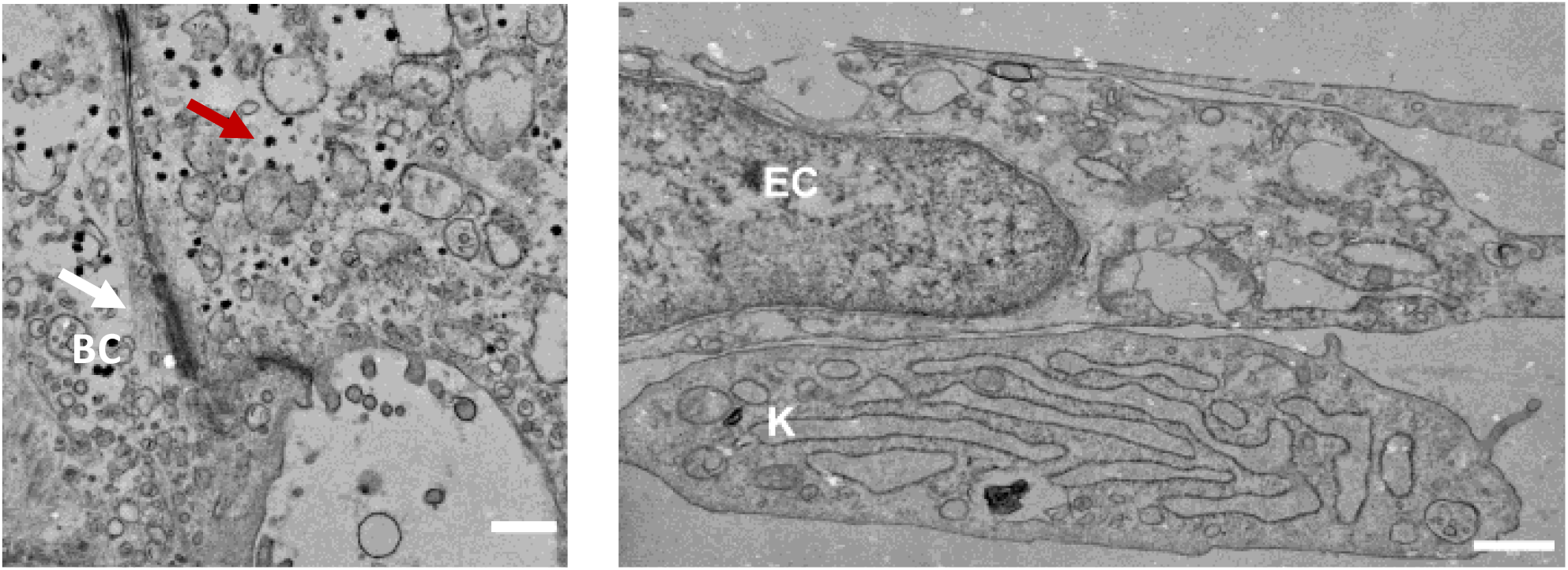
Representative transmission electron microscopy images showing a well-formed bile canaliculus (bc) between neighboring hepatocytes (left) and cell-cell contact formation between a Kupffer (K) cell and liver sinusoidal endothelial cell (right) (bar, 0.5 µm).

**Supplementary Figure S2.**
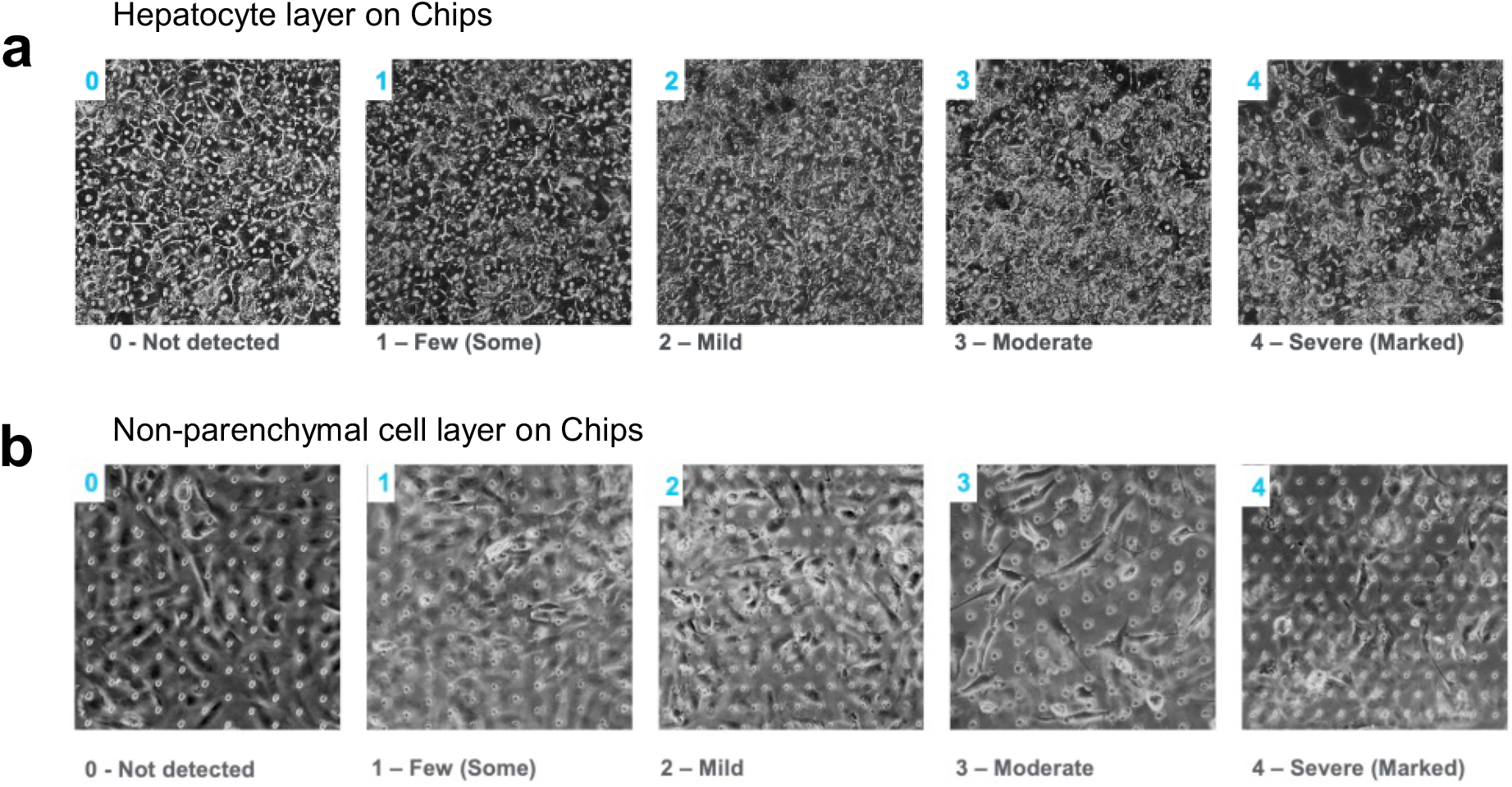
a) Representative brightfield images to depict the cellular morphology score in the top channel of the Liver-Chip which contains hepatocytes. A score of 0 represents no hepatotoxicity detected, which is defined by 95-100% healthy hepatocyte morphology, hexagonal shape containing binucleated cells, clear cell cytoplasm, distinctive cell junctions and less than 5% dead cells. A score of 1 represents at least 85% healthy hepatocyte morphology, a hexagonal shape containing binucleated cells, clear cell cytoplasm, distinctive cell junctions but < 15% are dead cells. A score of 2 represents mild hepatotoxicity with > 70% monolayer of hepatocytes visible, evidence that cells have begun to lose their distinct cell junctions, many cells contain a granulated cytoplasm but < 30% are dead cells. A score of 3 represents moderate hepatotoxicity with severe granulation of cytoplasm and most of the cells have lost their junctions. Approximately 50% of the cells are considered dead. A score of 4 represents severe hepatotoxicity with agglomeration of cell debris and > 50% of the cells are considered dead. The pores on the membrane become visible as there is no longer a cellular monolayer. b) Representative brightfield images to depict the cellular morphology score in the bottom channel of the Liver-Chip which contains non-parenchymal cells. A score of 0 represents no cytotoxicity detected, with an intact monolayer and <1% of the cells are dead. A score of 1 represents at least 90% of the monolayer is present and there are <10% dead cells. A score of 2 represents mild cytotoxicity with > 80% of the monolayer present and< 20% are dead cells. A score of 3 represents moderate cytotoxicity with > 50% of the monolayer present and < 50% are dead cells. A score of 4 represents severe cytotoxicity with > 50% of the cells are considered dead.

**Supplementary Figure S3.**
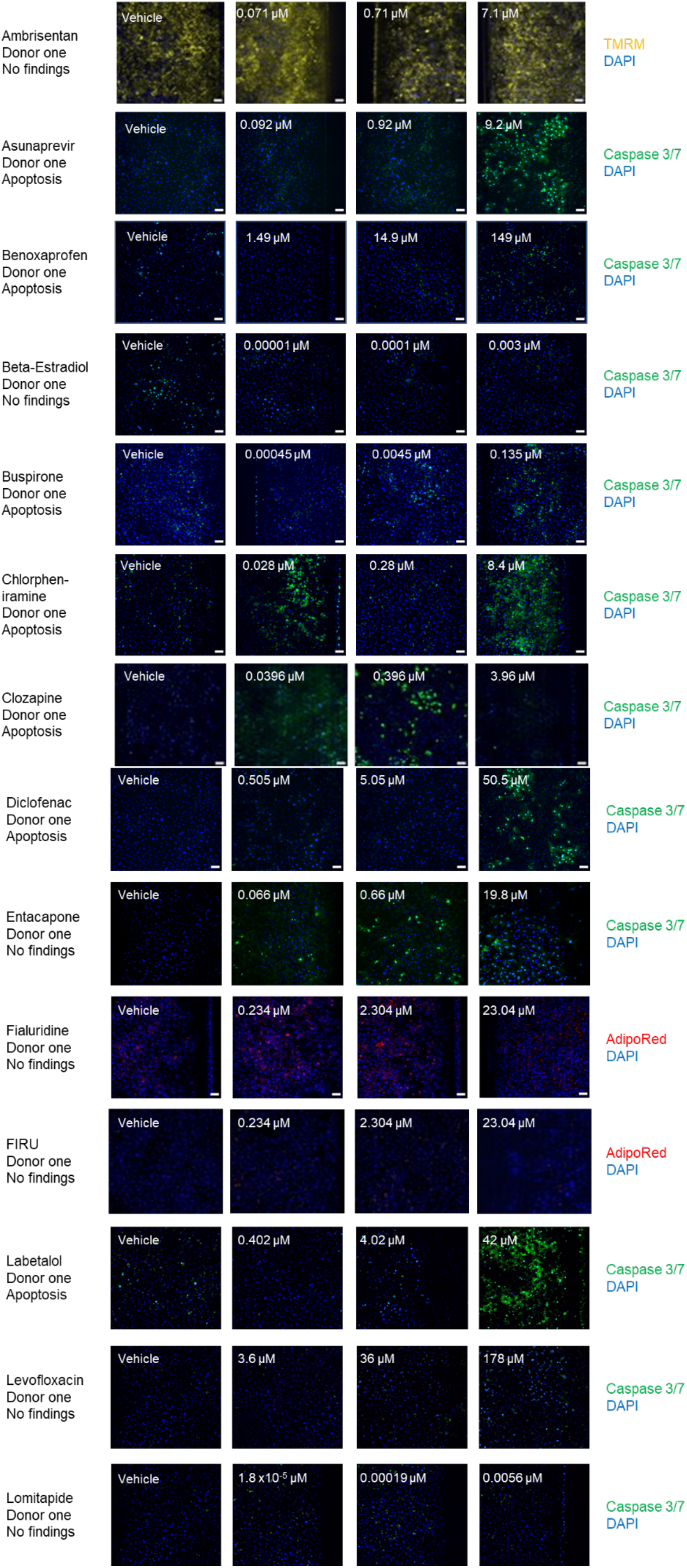
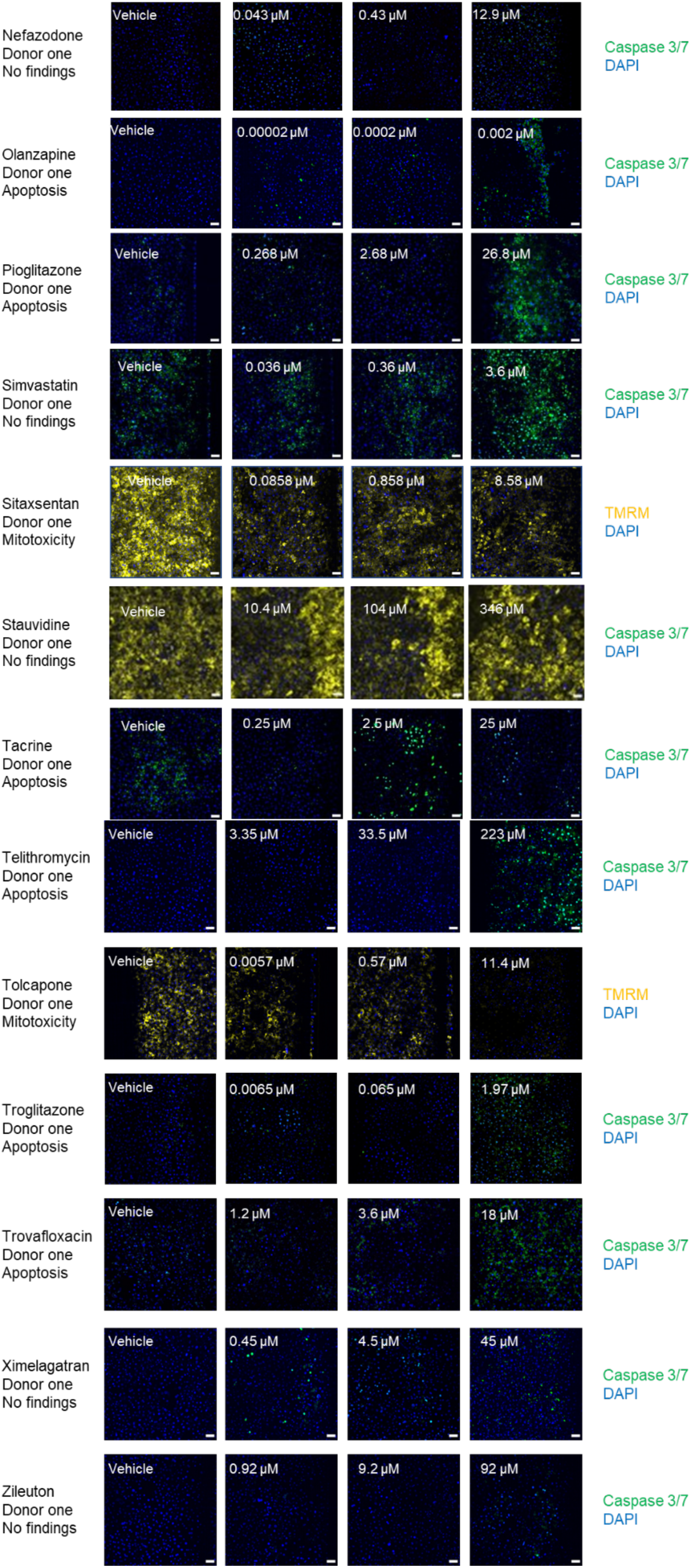
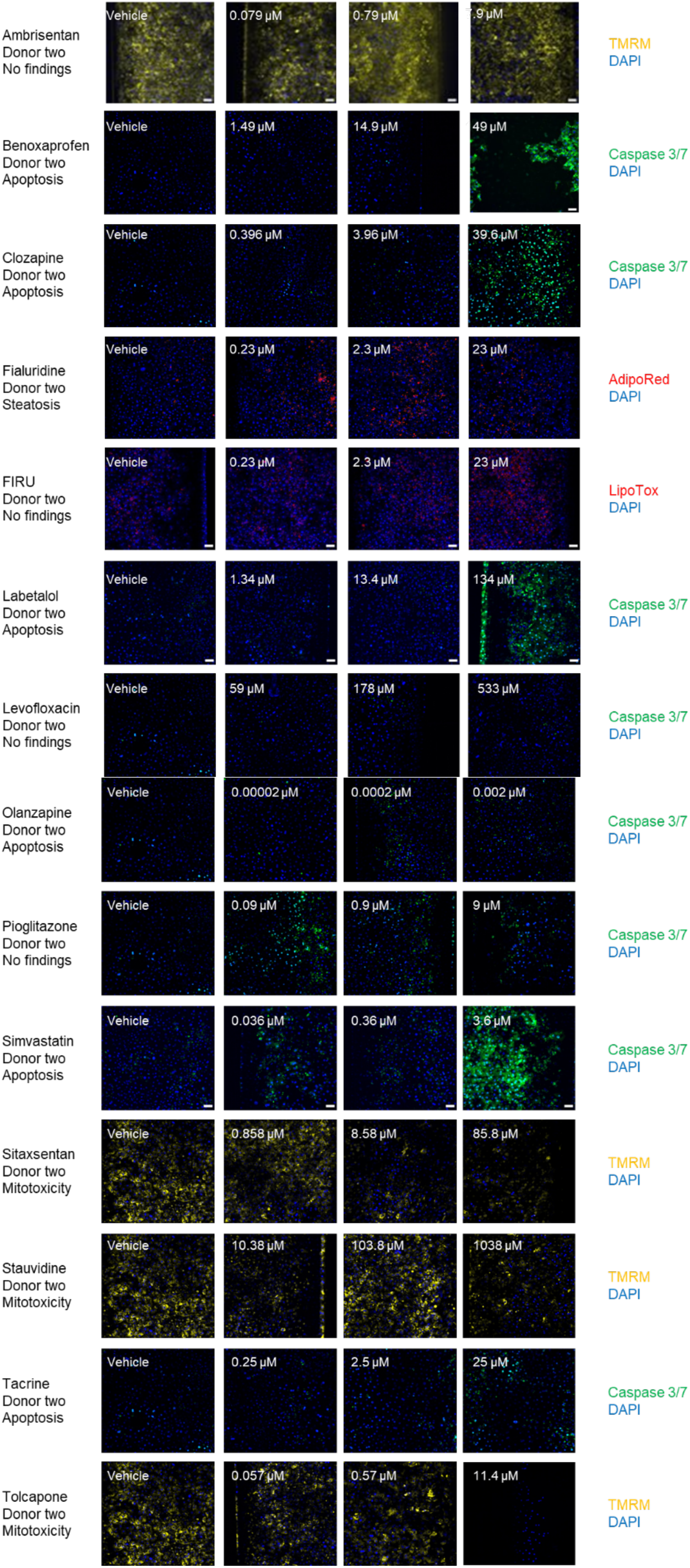
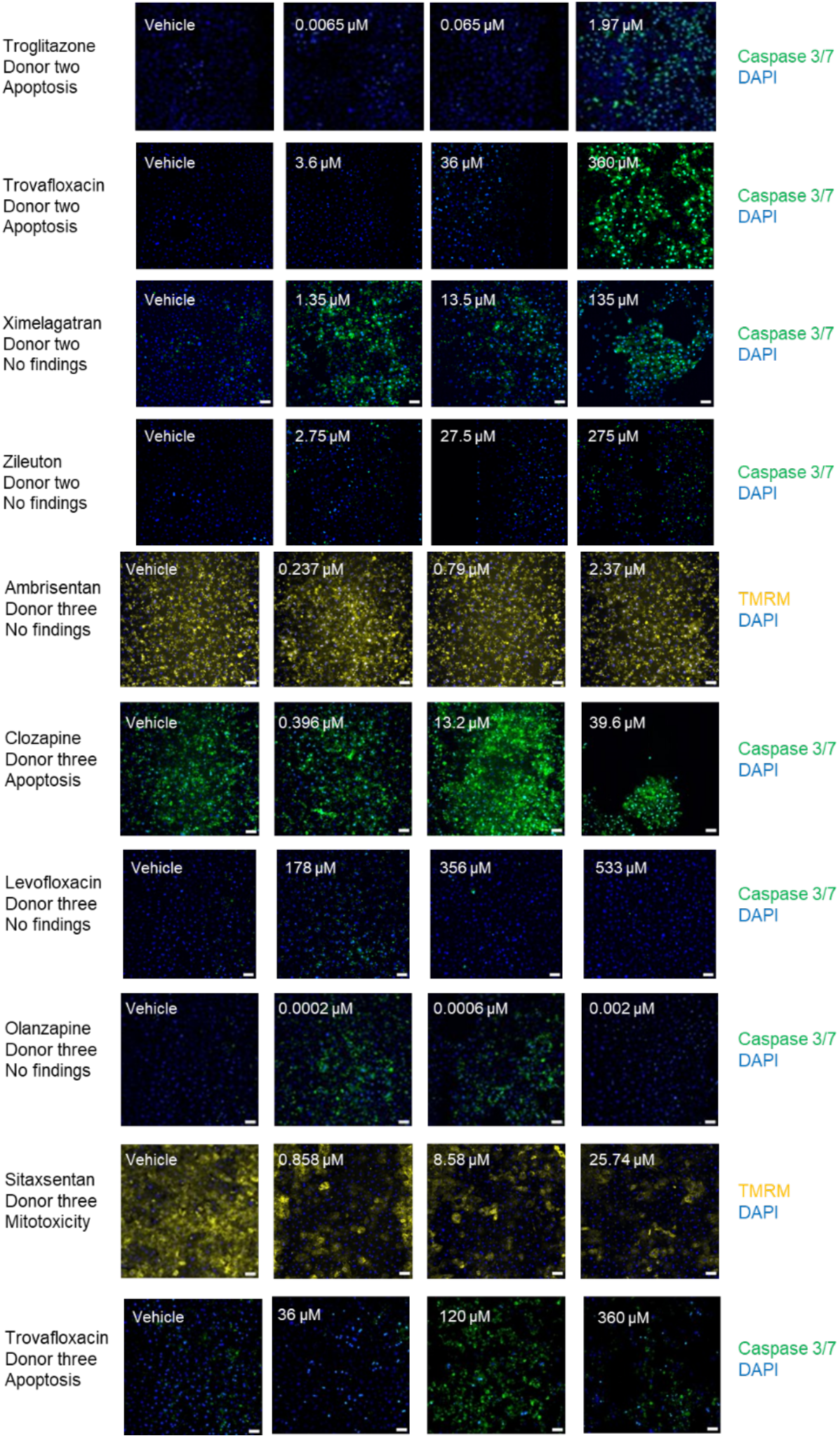
Representative immunofluorescent images from day 7 post-vehicle or drug administration of the hepatocyte cell layer in the top channel of the chip. Each drug is shown with its free drug concentration and corresponding vehicle image that was used for thresholding across each donor the drug was tested in. The data support the immunofluorescent findings statement in Table 2 and 3.

**Supplementary Table S1.**
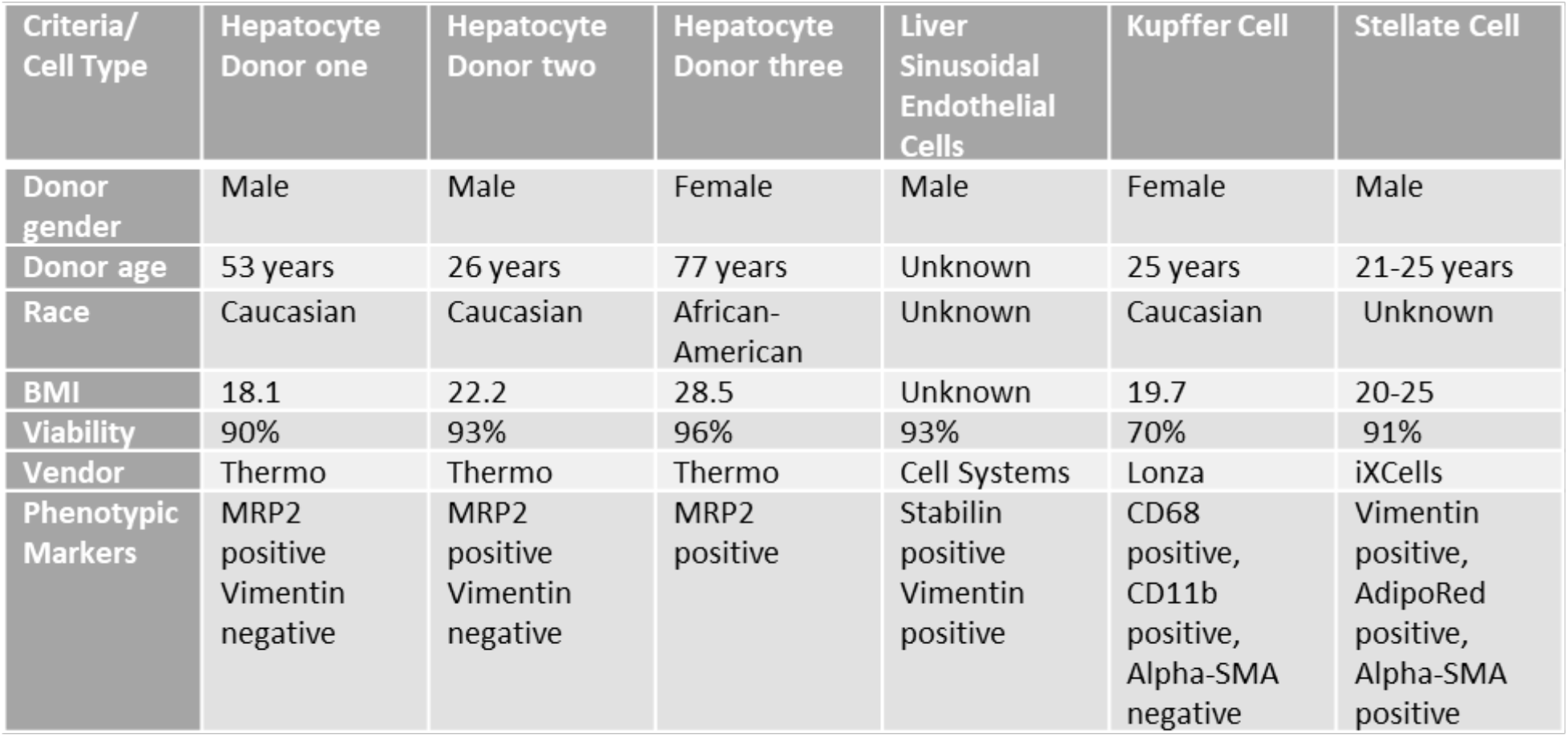
Details of the cell sources and their defining characteristics used in the investigation.

**Supplementary Table S2.**
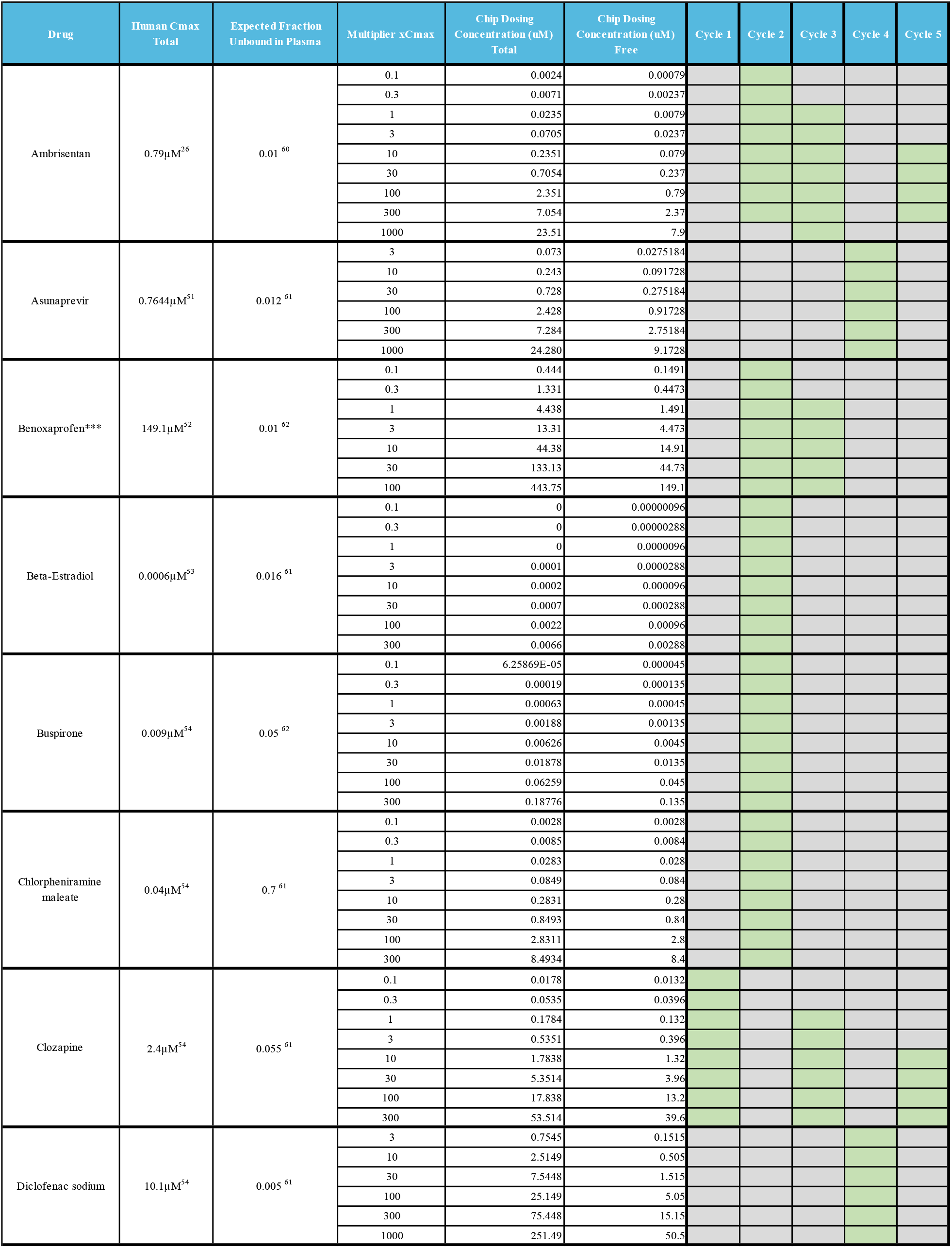
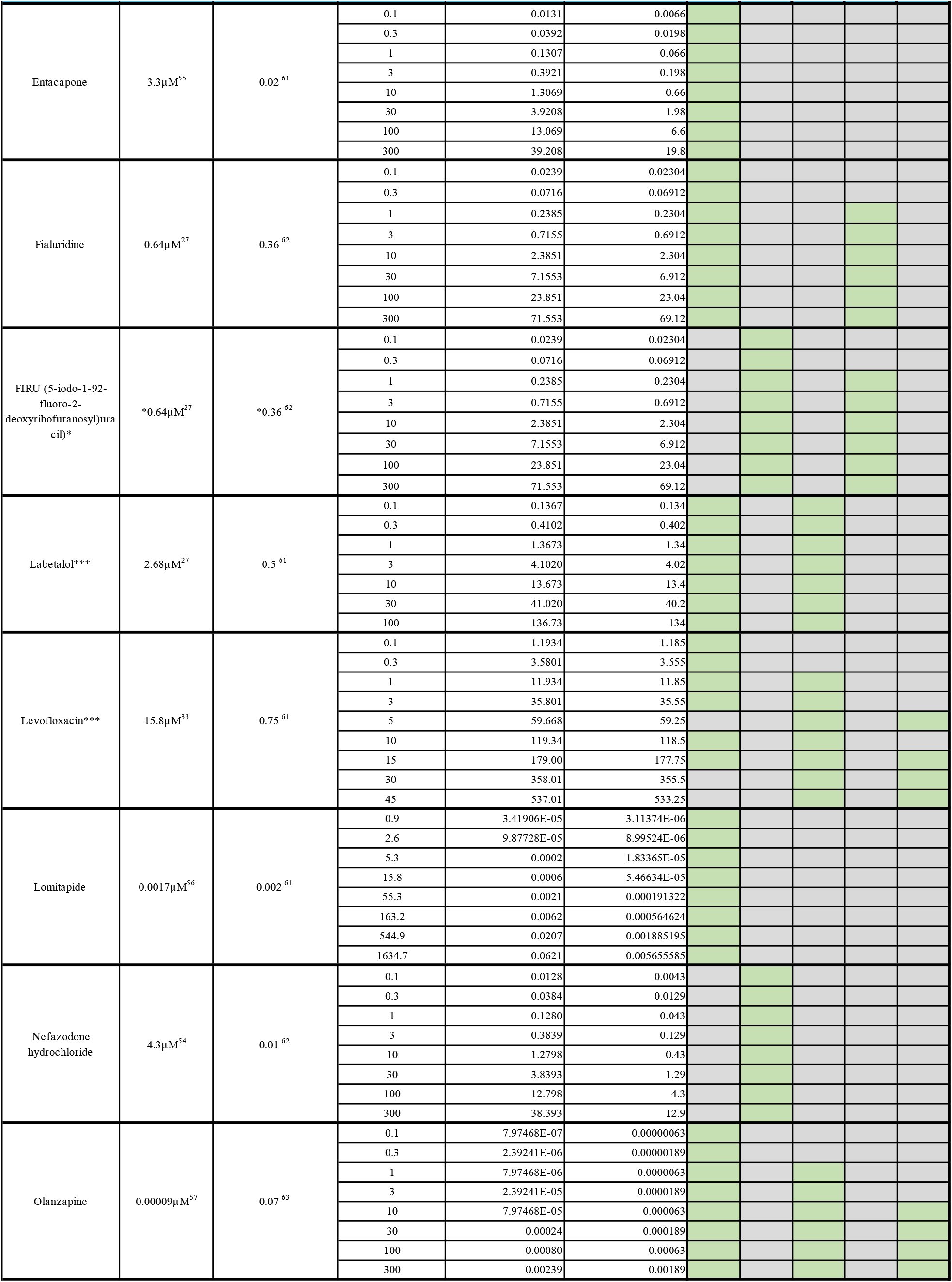
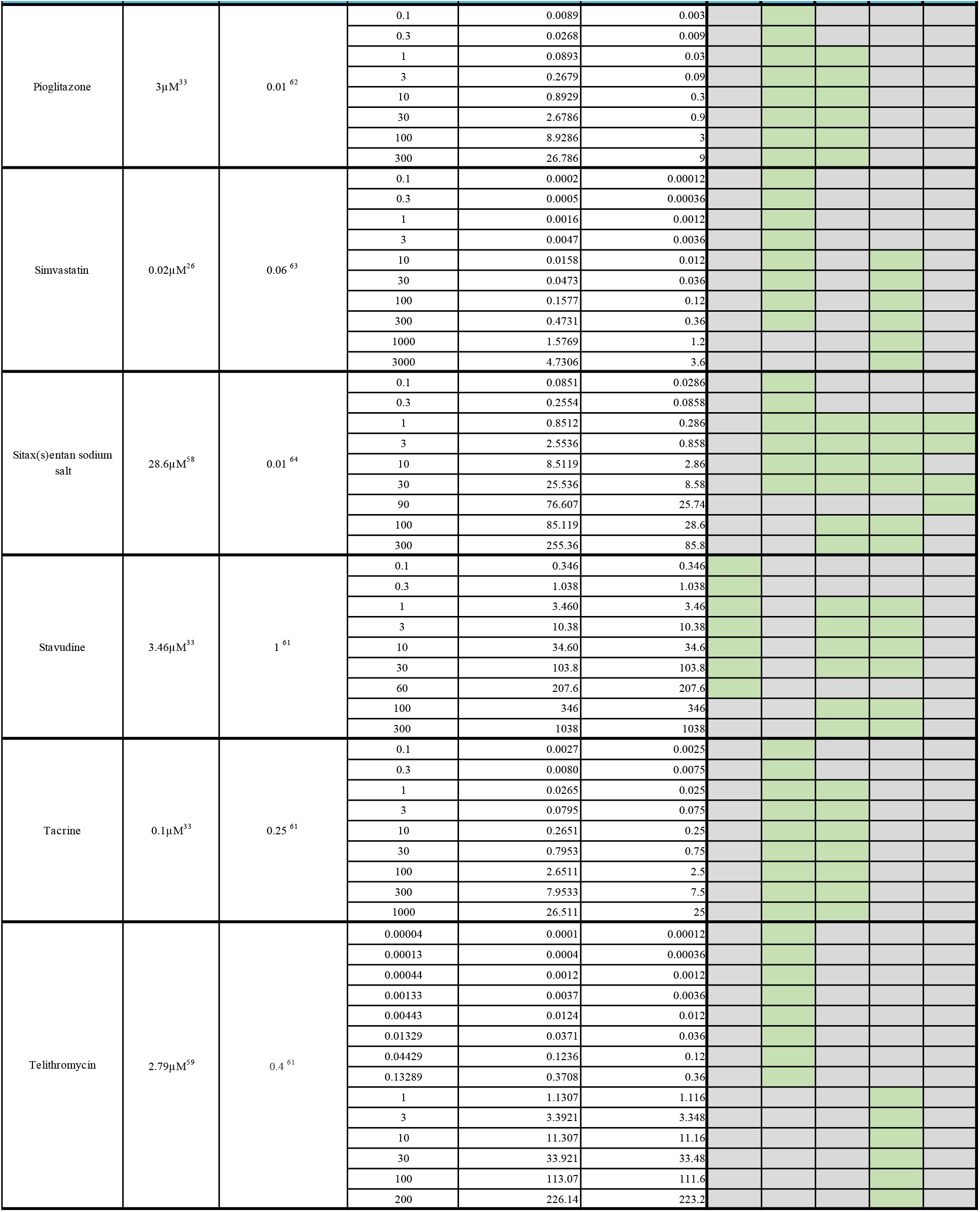
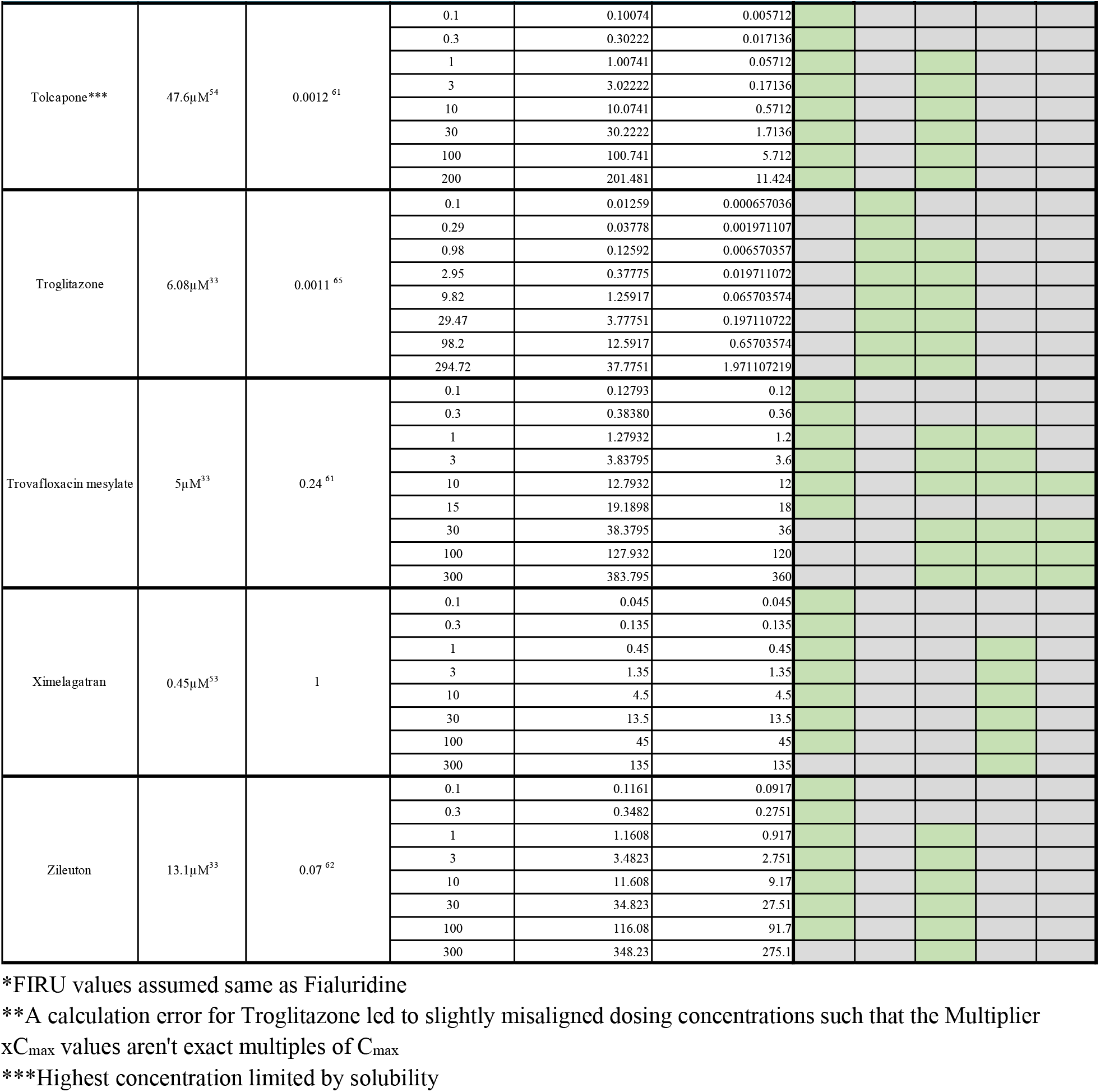
Drug information and dosing concentrations used in the investigation. Cycle-specific concentrations have been added for further clarity.

**Supplementary Table S3.**
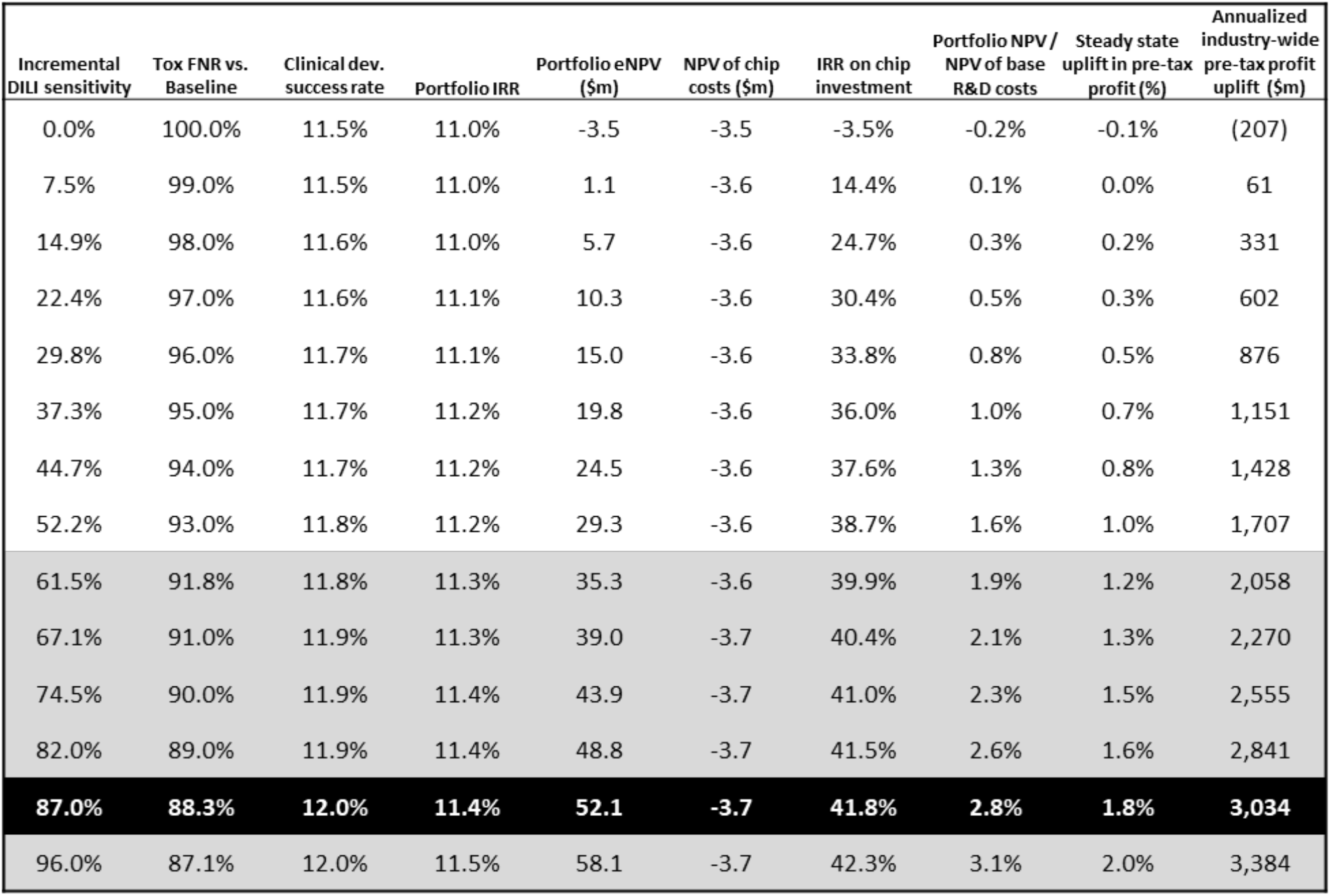
Portfolio value and industry profits increase with reductions in the false negative rate (FNR) of the preclinical toxicology assessment, which cause fewer toxic drugs to enter the clinic. The leftmost column tabulates the proportional improvement in DILI detection versus the base case, and the next column shows the improvement in the toxicology FNR relative to the model’s base case. Since DILI is around 13% or tox failure in development, near total DILI sensitivity can reduce the FNR to around 87% of its base case value. Subsequent columns then show the clinical development success rate from entry into Phase I to launch, the internal rate of return (IRR) of the R&D portfolio, the NPV of the portfolio discounted to the time of drug launch, the capitalized cost of Liver-Chips used in assessing the portfolio (discounted to the time of launch), and the marginal IRR on the Liver-Chip investment. The remaining columns capture the value uplift due to FNR improvement as a percentage uplift of the portfolio’s NPV relative to the baseline NPV of R&D, percentage uplift of steady state pre-tax profits, and the estimated increase in annual pre-tax profits for the small-molecule drug development industry. The row highlighted in dark gray relates to the improvement in FNR that may result from incorporating the Liver-Chip into DILI prediction workflows in accordance with the 87% sensitivity estimated by the present study. The rows highlighted in light gray correspond to the 95% confidence interval around this point estimate. The economic model behind these calculations is provided in full in the Supplementary Materials.

