## Supplemental tables and figures for "Qualifying a human Liver-Chip for predictive toxicology: Performance assessment and economic implications"

Ewart et al.

**Supplementary Material**

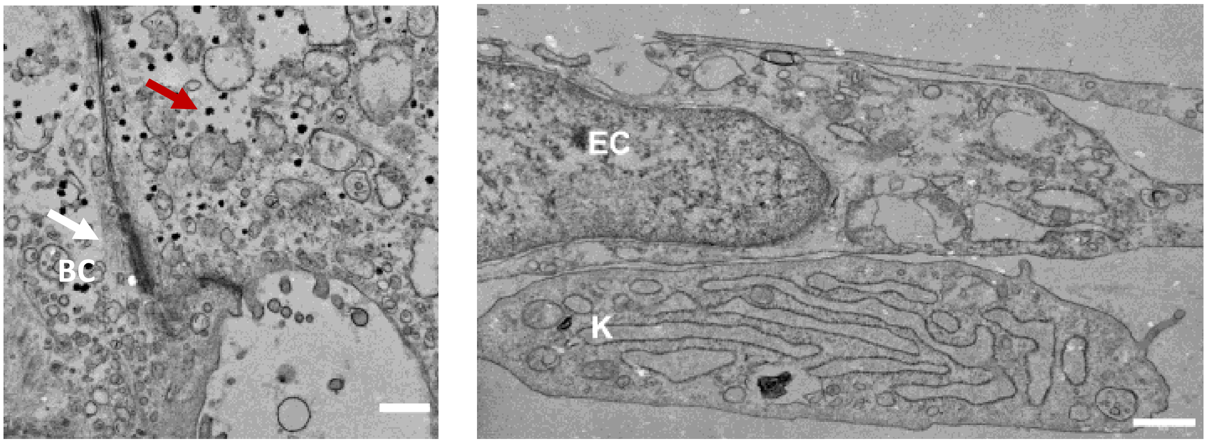


**Supplementary Figure S1.** Representative transmission electron microscopy images showing a well-formed bile canaliculus (bc) between neighboring hepatocytes (left) and cell-cell contact formation between a Kupffer (K) cell and liver sinusoidal endothelial cell (right) (bar, 0.5 µm).

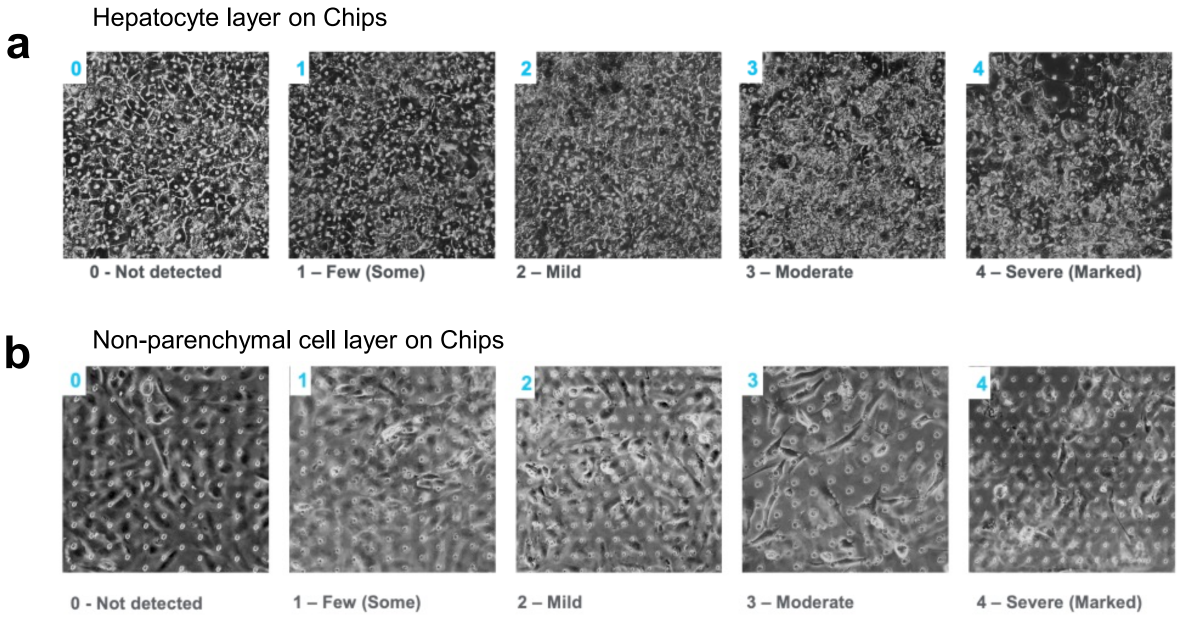


**Supplementary Figure S2.** a) Representative brightfield images to depict the cellular morphology score in the top channel of the Liver-Chip which contains hepatocytes. A score of 0 represents no hepatotoxicity detected, which is defined by 95-100% healthy hepatocyte morphology, hexagonal shape containing binucleated cells, clear cell cytoplasm, distinctive cell junctions and less than 5% dead cells. A score of 1 represents at least 85% healthy hepatocyte morphology, a hexagonal shape containing binucleated cells, clear cell cytoplasm, distinctive cell junctions but < 15% are dead cells. A score of 2 represents mild hepatotoxicity with > 70% monolayer of hepatocytes visible, evidence that cells have begun to lose their distinct cell junctions, many cells contain a granulated cytoplasm but < 30% are dead cells. A score of 3 represents moderate hepatotoxicity with severe granulation of cytoplasm and most of the cells have lost their junctions. Approximately 50% of the cells are considered dead. A score of 4 represents severe hepatotoxicity with agglomeration of cell debris and > 50% of the cells are considered dead. The pores on the membrane become visible as there is no longer a cellular monolayer. b) Representative brightfield images to depict the cellular morphology score in the bottom channel of the Liver-Chip which contains non-parenchymal cells. A score of 0 represents no cytotoxicity detected, with an intact monolayer and <1% of the cells are dead. A score of 1 represents at least 90% of the monolayer is present and there are <10% dead cells. A score of 2 represents mild cytotoxicity with > 80% of the monolayer present and < 20% are dead cells. A score of 3 represents moderate cytotoxicity with > 50% of the monolayer present and < 50% are dead cells. A score of 4 represents severe cytotoxicity with > 50% of the cells are considered dead.

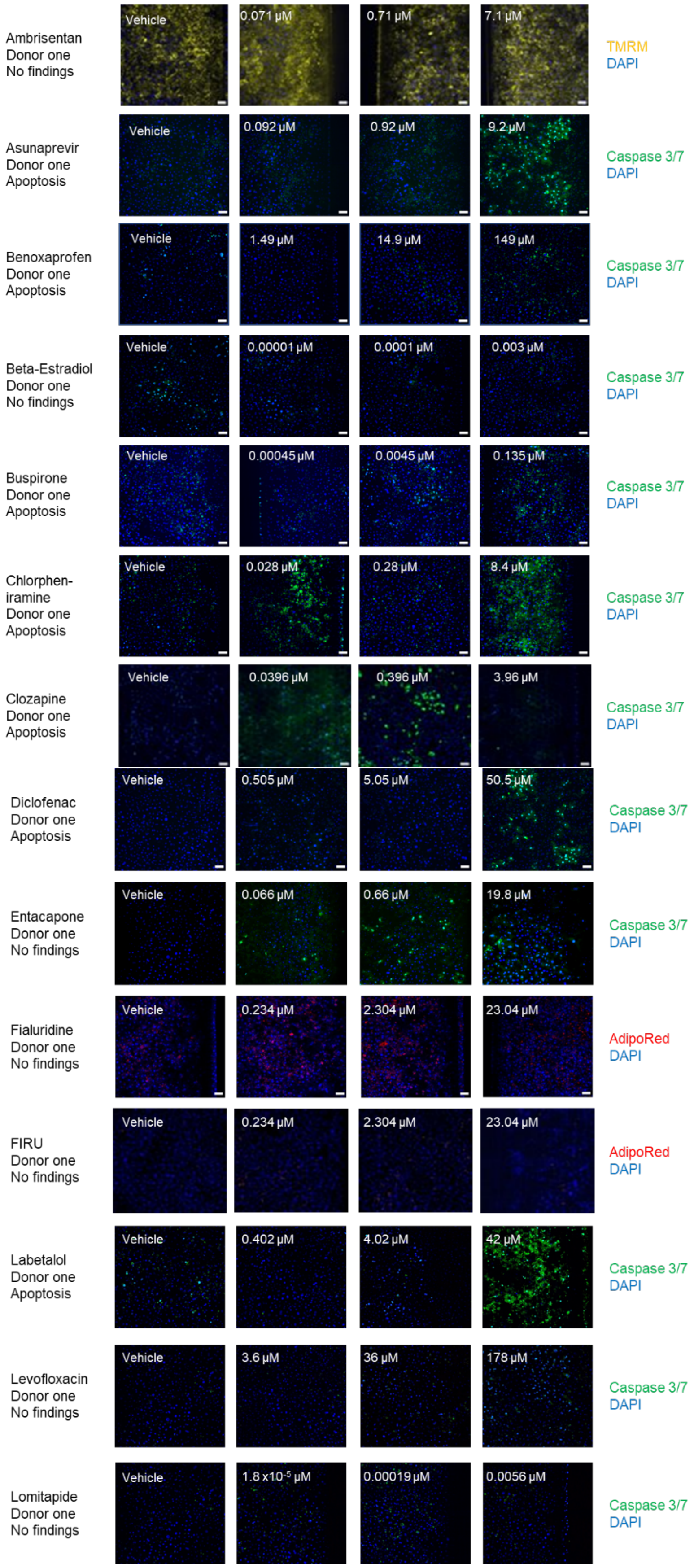

Ambrisentan
Donor one
No findings
Vehicle
0.071 µM
0.71 µM
7.1 µM
TMRM
DAPI
Asunaprevir
Donor one
Apoptosis
Vehicle
0.092 µM
0.92 µM
9.2 µM
Caspase 3/7
DAPI
Benoxaprofen
Donor one
Apoptosis
Vehicle
1.49 µM
14.9 µM
149 µM
Caspase 3/7
DAPI
Beta-Estradiol
Donor one
No findings
Vehicle
0.00001 µM
0.0001 µM
0.003 µM
Caspase 3/7
DAPI
Buspirone
Donor one
Apoptosis
Vehicle
0.00045 µM
0.0045 µM
0.135 µM
Caspase 3/7
DAPI
Chlorphen-
iramine
Donor one
Apoptosis
Vehicle
0.028 µM
0.28 µM
8.4 µM
Caspase 3/7
DAPI
Clozapine
Donor one
Apoptosis
Vehicle
0.0396 µM
0.396 µM
3.96 µM
Caspase 3/7
DAPI
Diclofenac
Donor one
Apoptosis
Vehicle
0.505 µM
5.05 µM
50.5 µM
Caspase 3/7
DAPI
Entacapone
Donor one
No findings
Vehicle
0.066 µM
0.66 µM
19.8 µM
Caspase 3/7
DAPI
Fialuridine
Donor one
No findings
Vehicle
0.234 µM
2.304 µM
23.04 µM
AdipoRed
DAPI
FIRU
Donor one
No findings
Vehicle
0.234 µM
2.304 µM
23.04 µM
AdipoRed
DAPI
Labetalol
Donor one
Apoptosis
Vehicle
0.402 µM
4.02 µM
42 µM
Caspase 3/7
DAPI
Levofloxacin
Donor one
No findings
Vehicle
3.6 µM
36 µM
178 µM
Caspase 3/7
DAPI
Lomitapide
Donor one
No findings
Vehicle
$1.8 x10^{-5}$ µM
0.00019 µM
0.0056 µM
Caspase 3/7
DAPI

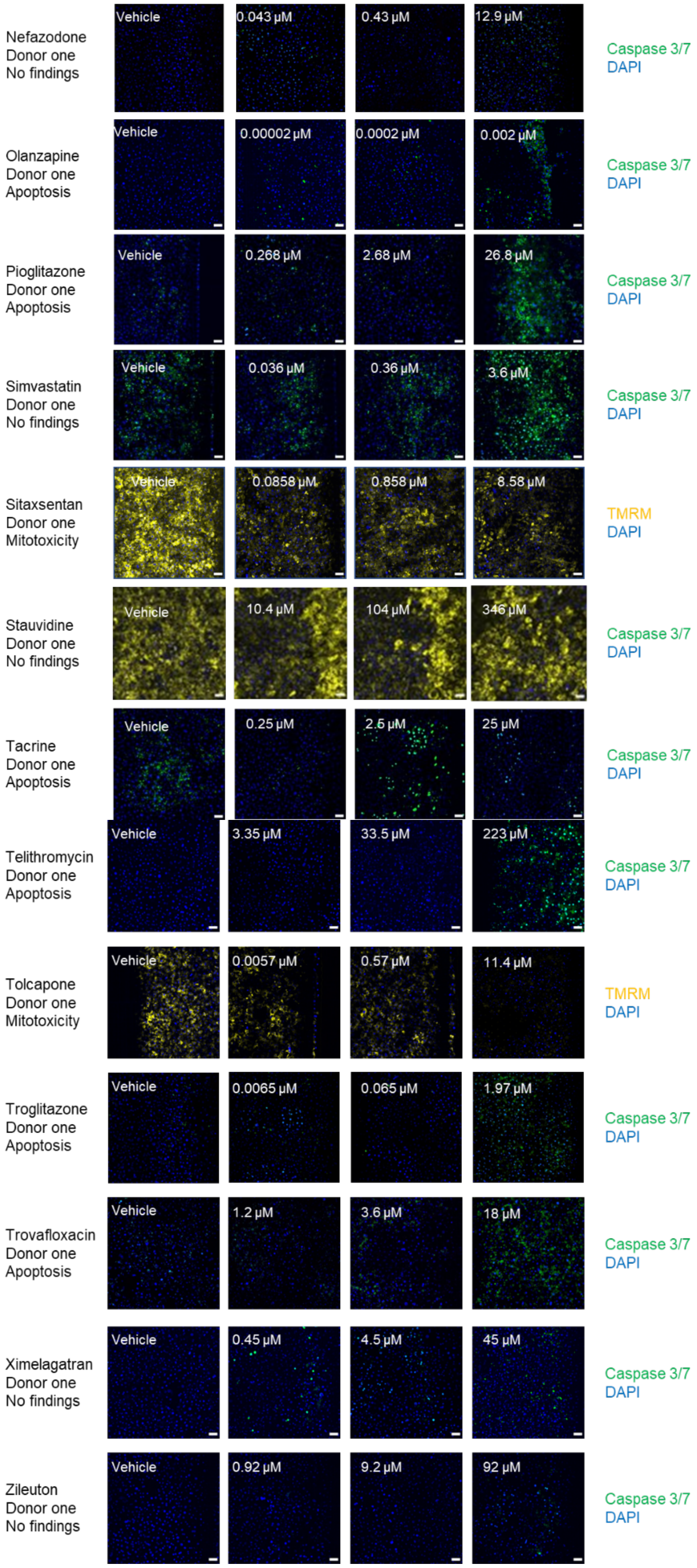
Nefazodone
Donor one
No findings
Vehicle
0.043 µM
0.43 µM
12.9 µM
Caspase 3/7
DAPI
Olanzapine
Donor one
Apoptosis
Vehicle
0.00002 µM
0.0002 µM
0.002 µM
Caspase 3/7
DAPI
Pioglitazone
Donor one
Apoptosis
Vehicle
0.268 µM
2.68 µM
26.8 µM
Caspase 3/7
DAPI
Simvastatin
Donor one
No findings
Vehicle
0.036 µM
0.36 µM
3.6 µM
Caspase 3/7
DAPI
Sitaxsentan
Donor one
Mitotoxicity
Vehicle
0.0858 µM
0.858 µM
8.58 µM
TMRM
DAPI
Stauvidine
Donor one
No findings
Vehicle
10.4 µM
104 µM
346 µM
Caspase 3/7
DAPI
Tacrine
Donor one
Apoptosis
Vehicle
0.25 µM
2.5 µM
25 µM
Caspase 3/7
DAPI
Telithromycin
Donor one
Apoptosis
Vehicle
3.35 µM
33.5 µM
223 µM
Caspase 3/7
DAPI
Tolcapone
Donor one
Mitotoxicity
Vehicle
0.0057 µM
0.57 µM
11.4 µM
TMRM
DAPI
Troglitazone
Donor one
Apoptosis
Vehicle
0.0065 µM
0.065 µM
1.97 µM
Caspase 3/7
DAPI
Trovafloxacin
Donor one
Apoptosis
Vehicle
1.2 µM
3.6 µM
18 µM
Caspase 3/7
DAPI
Ximelagatran
Donor one
No findings
Vehicle
0.45 µM
4.5 µM
45 µM
Caspase 3/7
DAPI
Zileuton
Donor one
No findings
Vehicle
0.92 µM
9.2 µM
92 µM
Caspase 3/7
DAPI

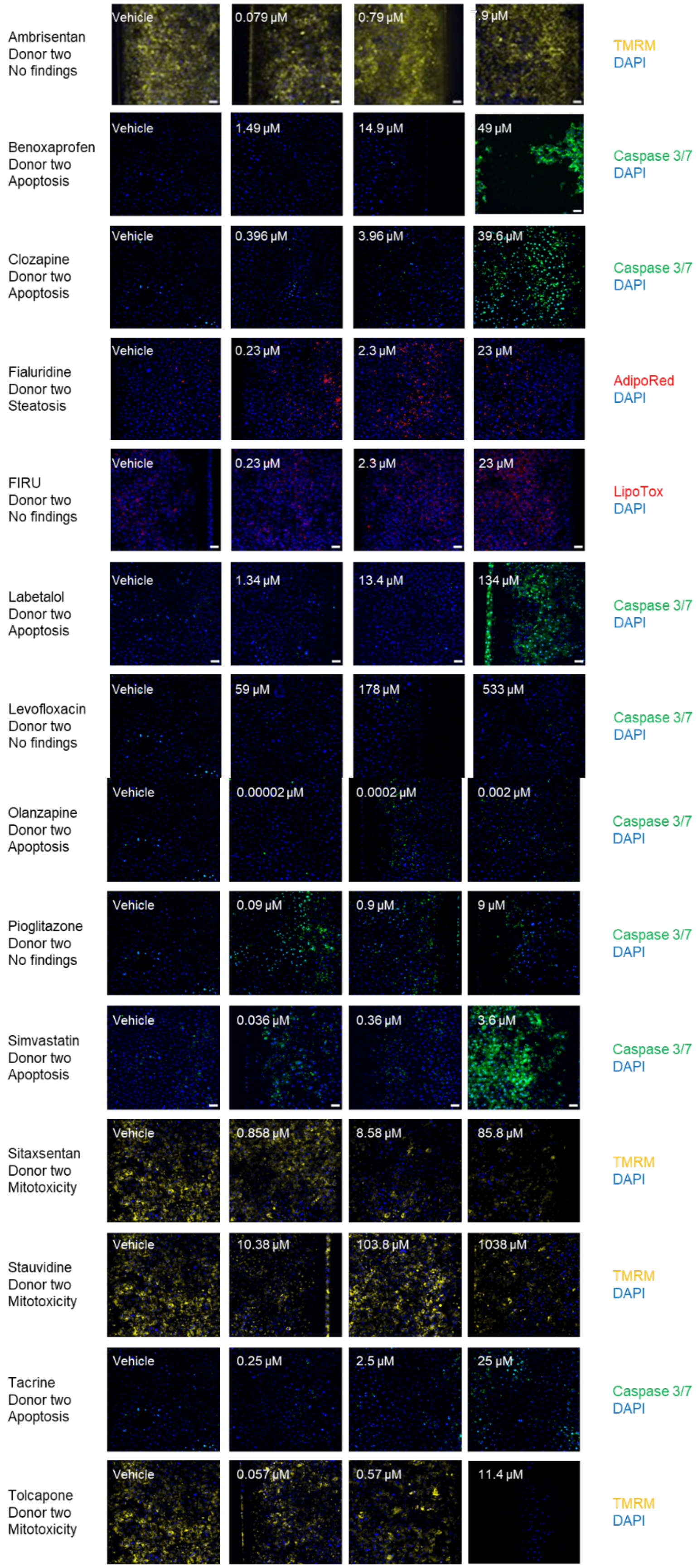
Ambrisentan
Donor two
No findings
Vehicle
0.079 µM
0.79 µM
9 µM
TMRM
DAPI
Benoxaprofen
Donor two
Apoptosis
Vehicle
1.49 µM
14.9 µM
49 µM
Caspase 3/7
DAPI
Clozapine
Donor two
Apoptosis
Vehicle
0.396 µM
3.96 µM
39.6 µM
Caspase 3/7
DAPI
Fialuridine
Donor two
Steatosis
Vehicle
0.23 µM
2.3 µM
23 µM
AdipoRed
DAPI
FIRU
Donor two
No findings
Vehicle
0.23 µM
2.3 µM
23 µM
LipoTox
DAPI
Labetalol
Donor two
Apoptosis
Vehicle
1.34 µM
13.4 µM
134 µM
Caspase 3/7
DAPI
Levofloxacin
Donor two
No findings
Vehicle
59 µM
178 µM
533 µM
Caspase 3/7
DAPI
Olanzapine
Donor two
Apoptosis
Vehicle
0.00002 µM
0.0002 µM
0.002 µM
Caspase 3/7
DAPI
Pioglitazone
Donor two
No findings
Vehicle
0.09 µM
0.9 µM
9 µM
Caspase 3/7
DAPI
Simvastatin
Donor two
Apoptosis
Vehicle
0.036 µM
0.36 µM
3.6 µM
Caspase 3/7
DAPI
Sitaxsentan
Donor two
Mitotoxicity
Vehicle
0.858 µM
8.58 µM
85.8 µM
TMRM
DAPI
Stauvidine
Donor two
Mitotoxicity
Vehicle
10.38 µM
103.8 µM
1038 µM
TMRM
DAPI
Tacrine
Donor two
Apoptosis
Vehicle
0.25 µM
2.5 µM
25 µM
Caspase 3/7
DAPI
Tolcapone
Donor two
Mitotoxicity
Vehicle
0.057 µM
0.57 µM
11.4 µM
TMRM
DAPI

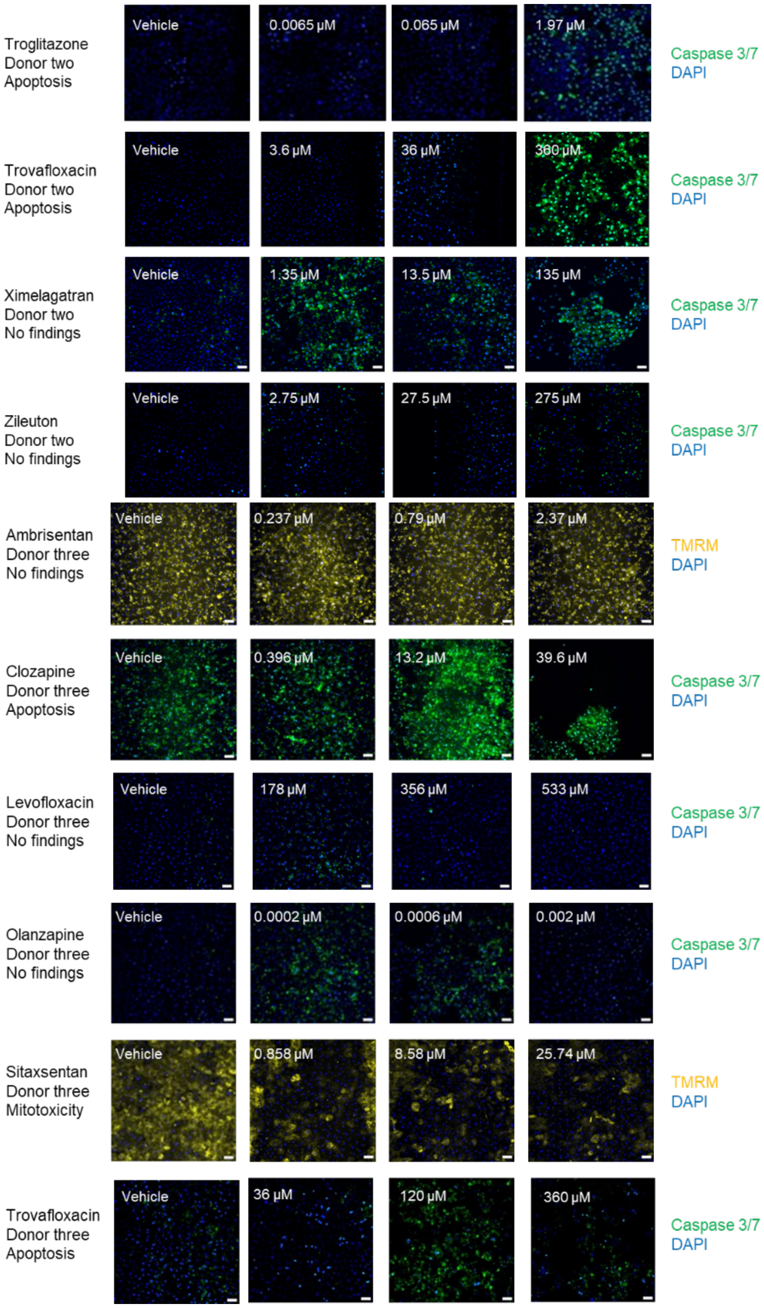


**Supplementary Figure S3.** Representative immunofluorescent images from day 7 post-vehicle or drug administration of the hepatocyte cell layer in the top channel of the chip. Each drug is shown with its free drug concentration and corresponding vehicle image that was used for thresholding across each donor the drug was tested in. The data support the immunofluorescent findings statement in Table 2 and 3.

| Criteria/ Cell Type | Hepatocyte Donor one | Hepatocyte Donor two | Hepatocyte Donor three | Liver Sinusoidal Endothelial Cells | Kupffer Cell | Stellate Cell |
|---|---|---|---|---|---|---|
| Donor gender | Male | Male | Female | Male | Female | Male |
| Donor age | 53 years | 26 years | 77 years | Unknown | 25 years | 21-25 years |
| Race | Caucasian | Caucasian | African-American | Unknown | Caucasian | Unknown |
| BMI | 18.1 | 22.2 | 28.5 | Unknown | 19.7 | 20-25 |
| Viability | 90% | 93% | 96% | 93% | 70% | 91% |
| Vendor | Thermo | Thermo | Thermo | Cell Systems | Lonza | iXCells |
| Phenotypic Markers | MRP2 positive Vimentin negative | MRP2 positive Vimentin negative | MRP2 positive | Stabilin positive Vimentin positive | CD68 positive, CD11b positive, Alpha-SMA negative | Vimentin positive, AdipoRed positive, Alpha-SMA positive |

**Supplementary Table S1.** Details of the cell sources and their defining characteristics used in the investigation.

| Drug | Human Cmax Total | Expected Fraction Unbound in Plasma | Multiplier xCmax | Chip Dosing Concentration (uM) Total | Chip Dosing Concentration (uM) Free | Cycle 1 | Cycle 2 | Cycle 3 | Cycle 4 | Cycle 5 |
|---|---|---|---|---|---|---|---|---|---|---|
| Ambrisentan | 0.79µM[26] | 0.01 [60] | 0.1 | 0.0024 | 0.00079 | | | | | |
| | | | 0.3 | 0.0071 | 0.00237 | | | | | |
| | | | 1 | 0.0235 | 0.0079 | | | | | |
| | | | 3 | 0.0705 | 0.0237 | | | | | |
| | | | 10 | 0.2351 | 0.079 | | | | | |
| | | | 30 | 0.7054 | 0.237 | | | | | |
| | | | 100 | 2.351 | 0.79 | | | | | |
| | | | 300 | 7.054 | 2.37 | | | | | |
| | | | 1000 | 23.51 | 7.9 | | | | | |
| Asunaprevir | 0.7644µM[51] | 0.012 [61] | 3 | 0.073 | 0.0275184 | | | | | |
| | | | 10 | 0.243 | 0.091728 | | | | | |
| | | | 30 | 0.728 | 0.275184 | | | | | |
| | | | 100 | 2.428 | 0.91728 | | | | | |
| | | | 300 | 7.284 | 2.75184 | | | | | |
| | | | 1000 | 24.280 | 9.1728 | | | | | |
| Benoxaprofen*** | 149.1µM[52] | 0.01 [62] | 0.1 | 0.444 | 0.1491 | | | | | |
| | | | 0.3 | 1.331 | 0.4473 | | | | | |
| | | | 1 | 4.438 | 1.491 | | | | | |
| | | | 3 | 13.31 | 4.473 | | | | | |
| | | | 10 | 44.38 | 14.91 | | | | | |
| | | | 30 | 133.13 | 44.73 | | | | | |
| | | | 100 | 443.75 | 149.1 | | | | | |
| Beta-Estradiol | 0.0006µM[53] | 0.016 [61] | 0.1 | 0 | 0.00000096 | | | | | |
| | | | 0.3 | 0 | 0.00000288 | | | | | |
| | | | 1 | 0 | 0.0000096 | | | | | |
| | | | 3 | 0.0001 | 0.0000288 | | | | | |
| | | | 10 | 0.0002 | 0.000096 | | | | | |
| | | | 30 | 0.0007 | 0.000288 | | | | | |
| | | | 100 | 0.0022 | 0.00096 | | | | | |
| | | | 300 | 0.0066 | 0.00288 | | | | | |
| Buspirone | 0.009µM[54] | 0.05 [62] | 0.1 | 6.25869E-05 | 0.000045 | | | | | |
| | | | 0.3 | 0.00019 | 0.000135 | | | | | |
| | | | 1 | 0.00063 | 0.00045 | | | | | |
| | | | 3 | 0.00188 | 0.00135 | | | | | |
| | | | 10 | 0.00626 | 0.0045 | | | | | |
| | | | 30 | 0.01878 | 0.0135 | | | | | |
| | | | 100 | 0.06259 | 0.045 | | | | | |
| | | | 300 | 0.18776 | 0.135 | | | | | |
| Chlorpheniramine maleate | 0.04µM[54] | 0.7 [61] | 0.1 | 0.0028 | 0.0028 | | | | | |
| | | | 0.3 | 0.0085 | 0.0084 | | | | | |
| | | | 1 | 0.0283 | 0.028 | | | | | |
| | | | 3 | 0.0849 | 0.084 | | | | | |
| | | | 10 | 0.2831 | 0.28 | | | | | |
| | | | 30 | 0.8493 | 0.84 | | | | | |
| | | | 100 | 2.8311 | 2.8 | | | | | |
| | | | 300 | 8.4934 | 8.4 | | | | | |
| Clozapine | 2.4µM[54] | 0.055 [61] | 0.1 | 0.0178 | 0.0132 | | | | | |
| | | | 0.3 | 0.0535 | 0.0396 | | | | | |
| | | | 1 | 0.1784 | 0.132 | | | | | |
| | | | 3 | 0.5351 | 0.396 | | | | | |
| | | | 10 | 1.7838 | 1.32 | | | | | |
| | | | 30 | 5.3514 | 3.96 | | | | | |
| | | | 100 | 17.838 | 13.2 | | | | | |
| | | | 300 | 53.514 | 39.6 | | | | | |
| Diclofenac sodium | 10.1µM[54] | 0.005 [61] | 3 | 0.7545 | 0.1515 | | | | | |
| | | | 10 | 2.5149 | 0.505 | | | | | |
| | | | 30 | 7.5448 | 1.515 | | | | | |
| | | | 100 | 25.149 | 5.05 | | | | | |
| | | | 300 | 75.448 | 15.15 | | | | | |
| | | | 1000 | 251.49 | 50.5 | | | | | |

| Drug | Human Cmax Total | Expected Fraction Unbound in Plasma | Multiplier xCmax | Chip Dosing Concentration (uM) Total | Chip Dosing Concentration (uM) Free | Cycle 1 | Cycle 2 | Cycle 3 | Cycle 4 | Cycle 5 |
|---|---|---|---|---|---|---|---|---|---|---|
| Entacapone | 3.3µM[55] | 0.02 [61] | 0.1 | 0.0131 | 0.0066 | | | | | |
| | | | 0.3 | 0.0392 | 0.0198 | | | | | |
| | | | 1 | 0.1307 | 0.066 | | | | | |
| | | | 3 | 0.3921 | 0.198 | | | | | |
| | | | 10 | 1.3069 | 0.66 | | | | | |
| | | | 30 | 3.9208 | 1.98 | | | | | |
| | | | 100 | 13.069 | 6.6 | | | | | |
| | | | 300 | 39.208 | 19.8 | | | | | |
| Fialuridine | 0.64µM[27] | 0.36 [62] | 0.1 | 0.0239 | 0.02304 | | | | | |
| | | | 0.3 | 0.0716 | 0.06912 | | | | | |
| | | | 1 | 0.2385 | 0.2304 | | | | | |
| | | | 3 | 0.7155 | 0.6912 | | | | | |
| | | | 10 | 2.3851 | 2.304 | | | | | |
| | | | 30 | 7.1553 | 6.912 | | | | | |
| | | | 100 | 23.851 | 23.04 | | | | | |
| | | | 300 | 71.553 | 69.12 | | | | | |
| FIRU (5-iodo-1-92-fluoro-2-deoxyribofuranosyl)uracil)* | *0.64µM[27] | *0.36 [62] | 0.1 | 0.0239 | 0.02304 | | | | | |
| | | | 0.3 | 0.0716 | 0.06912 | | | | | |
| | | | 1 | 0.2385 | 0.2304 | | | | | |
| | | | 3 | 0.7155 | 0.6912 | | | | | |
| | | | 10 | 2.3851 | 2.304 | | | | | |
| | | | 30 | 7.1553 | 6.912 | | | | | |
| | | | 100 | 23.851 | 23.04 | | | | | |
| | | | 300 | 71.553 | 69.12 | | | | | |
| Labetalol*** | 2.68µM[27] | 0.5 [61] | 0.1 | 0.1367 | 0.134 | | | | | |
| | | | 0.3 | 0.4102 | 0.402 | | | | | |
| | | | 1 | 1.3673 | 1.34 | | | | | |
| | | | 3 | 4.1020 | 4.02 | | | | | |
| | | | 10 | 13.673 | 13.4 | | | | | |
| | | | 30 | 41.020 | 40.2 | | | | | |
| | | | 100 | 136.73 | 134 | | | | | |
| Levofloxacin*** | 15.8µM[33] | 0.75 [61] | 0.1 | 1.1934 | 1.185 | | | | | |
| | | | 0.3 | 3.5801 | 3.555 | | | | | |
| | | | 1 | 11.934 | 11.85 | | | | | |
| | | | 3 | 35.801 | 35.55 | | | | | |
| | | | 5 | 59.668 | 59.25 | | | | | |
| | | | 10 | 119.34 | 118.5 | | | | | |
| | | | 15 | 179.00 | 177.75 | | | | | |
| | | | 30 | 358.01 | 355.5 | | | | | |
| | | | 45 | 537.01 | 533.25 | | | | | |
| Lomitapide | 0.0017µM[56] | 0.002 [61] | 0.9 | 3.41906E-05 | 3.11374E-06 | | | | | |
| | | | 2.6 | 9.87728E-05 | 8.99524E-06 | | | | | |
| | | | 5.3 | 0.0002 | 1.83365E-05 | | | | | |
| | | | 15.8 | 0.0006 | 5.46634E-05 | | | | | |
| | | | 55.3 | 0.0021 | 0.000191322 | | | | | |
| | | | 163.2 | 0.0062 | 0.000564624 | | | | | |
| | | | 544.9 | 0.0207 | 0.001885195 | | | | | |
| | | | 1634.7 | 0.0621 | 0.005655585 | | | | | |
| Nefazodone hydrochloride | 4.3µM[54] | 0.01 [62] | 0.1 | 0.0128 | 0.0043 | | | | | |
| | | | 0.3 | 0.0384 | 0.0129 | | | | | |
| | | | 1 | 0.1280 | 0.043 | | | | | |
| | | | 3 | 0.3839 | 0.129 | | | | | |
| | | | 10 | 1.2798 | 0.43 | | | | | |
| | | | 30 | 3.8393 | 1.29 | | | | | |
| | | | 100 | 12.798 | 4.3 | | | | | |
| | | | 300 | 38.393 | 12.9 | | | | | |
| Olanzapine | 0.00009µM[57] | 0.07 [63] | 0.1 | 7.97468E-07 | 0.00000063 | | | | | |
| | | | 0.3 | 2.39241E-06 | 0.00000189 | | | | | |
| | | | 1 | 7.97468E-06 | 0.0000063 | | | | | |
| | | | 3 | 2.39241E-05 | 0.0000189 | | | | | |
| | | | 10 | 7.97468E-05 | 0.000063 | | | | | |
| | | | 30 | 0.00024 | 0.000189 | | | | | |
| | | | 100 | 0.00080 | 0.00063 | | | | | |
| | | | 300 | 0.00239 | 0.00189 | | | | | |

| Drug | Human Cmax Total | Expected Fraction Unbound in Plasma | Multiplier xCmax | Chip Dosing Concentration (uM) Total | Chip Dosing Concentration (uM) Free | Cycle 1 | Cycle 2 | Cycle 3 | Cycle 4 | Cycle 5 |
|---|---|---|---|---|---|---|---|---|---|---|
| Pioglitazone | 3µM[33] | 0.01 [62] | 0.1 | 0.0089 | 0.003 | | | | | |
| | | | 0.3 | 0.0268 | 0.009 | | | | | |
| | | | 1 | 0.0893 | 0.03 | | | | | |
| | | | 3 | 0.2679 | 0.09 | | | | | |
| | | | 10 | 0.8929 | 0.3 | | | | | |
| | | | 30 | 2.6786 | 0.9 | | | | | |
| | | | 100 | 8.9286 | 3 | | | | | |
| | | | 300 | 26.786 | 9 | | | | | |
| Simvastatin | 0.02µM[26] | 0.06 [63] | 0.1 | 0.0002 | 0.00012 | | | | | |
| | | | 0.3 | 0.0005 | 0.00036 | | | | | |
| | | | 1 | 0.0016 | 0.0012 | | | | | |
| | | | 3 | 0.0047 | 0.0036 | | | | | |
| | | | 10 | 0.0158 | 0.012 | | | | | |
| | | | 30 | 0.0473 | 0.036 | | | | | |
| | | | 100 | 0.1577 | 0.12 | | | | | |
| | | | 300 | 0.4731 | 0.36 | | | | | |
| | | | 1000 | 1.5769 | 1.2 | | | | | |
| | | | 3000 | 4.7306 | 3.6 | | | | | |
| Sitax(s)entan sodium salt | 28.6µM[58] | 0.01 [64] | 0.1 | 0.0851 | 0.0286 | | | | | |
| | | | 0.3 | 0.2554 | 0.0858 | | | | | |
| | | | 1 | 0.8512 | 0.286 | | | | | |
| | | | 3 | 2.5536 | 0.858 | | | | | |
| | | | 10 | 8.5119 | 2.86 | | | | | |
| | | | 30 | 25.536 | 8.58 | | | | | |
| | | | 90 | 76.607 | 25.74 | | | | | |
| | | | 100 | 85.119 | 28.6 | | | | | |
| | | | 300 | 255.36 | 85.8 | | | | | |
| Stavudine | 3.46µM[33] | 1 [61] | 0.1 | 0.346 | 0.346 | | | | | |
| | | | 0.3 | 1.038 | 1.038 | | | | | |
| | | | 1 | 3.460 | 3.46 | | | | | |
| | | | 3 | 10.38 | 10.38 | | | | | |
| | | | 10 | 34.60 | 34.6 | | | | | |
| | | | 30 | 103.8 | 103.8 | | | | | |
| | | | 60 | 207.6 | 207.6 | | | | | |
| | | | 100 | 346 | 346 | | | | | |
| | | | 300 | 1038 | 1038 | | | | | |
| Tacrine | 0.1µM[33] | 0.25 [61] | 0.1 | 0.0027 | 0.0025 | | | | | |
| | | | 0.3 | 0.0080 | 0.0075 | | | | | |
| | | | 1 | 0.0265 | 0.025 | | | | | |
| | | | 3 | 0.0795 | 0.075 | | | | | |
| | | | 10 | 0.2651 | 0.25 | | | | | |
| | | | 30 | 0.7953 | 0.75 | | | | | |
| | | | 100 | 2.6511 | 2.5 | | | | | |
| | | | 300 | 7.9533 | 7.5 | | | | | |
| | | | 1000 | 26.511 | 25 | | | | | |
| Telithromycin | 2.79µM[59] | 0.4 [61] | 0.00004 | 0.0001 | 0.00012 | | | | | |
| | | | 0.00013 | 0.0004 | 0.00036 | | | | | |
| | | | 0.00044 | 0.0012 | 0.0012 | | | | | |
| | | | 0.00133 | 0.0037 | 0.0036 | | | | | |
| | | | 0.00443 | 0.0124 | 0.012 | | | | | |
| | | | 0.01329 | 0.0371 | 0.036 | | | | | |
| | | | 0.04429 | 0.1236 | 0.12 | | | | | |
| | | | 0.13289 | 0.3708 | 0.36 | | | | | |
| | | | 1 | 1.1307 | 1.116 | | | | | |
| | | | 3 | 3.3921 | 3.348 | | | | | |
| | | | 10 | 11.307 | 11.16 | | | | | |
| | | | 30 | 33.921 | 33.48 | | | | | |
| | | | 100 | 113.07 | 111.6 | | | | | |
| | | | 200 | 226.14 | 223.2 | | | | | |

| Drug | Human Cmax Total | Expected Fraction Unbound in Plasma | Multiplier xCmax | Chip Dosing Concentration (uM) Total | Chip Dosing Concentration (uM) Free | Cycle 1 | Cycle 2 | Cycle 3 | Cycle 4 | Cycle 5 |
|---|---|---|---|---|---|---|---|---|---|---|
| Tolcapone*** | 47.6μM[54] | 0.0012 [61] | 0.1 | 0.10074 | 0.005712 | | | | | |
| | | | 0.3 | 0.30222 | 0.017136 | | | | | |
| | | | 1 | 1.00741 | 0.05712 | | | | | |
| | | | 3 | 3.02222 | 0.17136 | | | | | |
| | | | 10 | 10.0741 | 0.5712 | | | | | |
| | | | 30 | 30.2222 | 1.7136 | | | | | |
| | | | 100 | 100.741 | 5.712 | | | | | |
| | | | 200 | 201.481 | 11.424 | | | | | |
| Troglitazone | 6.08μM[33] | 0.0011 [65] | 0.1 | 0.01259 | 0.000657036 | | | | | |
| | | | 0.29 | 0.03778 | 0.001971107 | | | | | |
| | | | 0.98 | 0.12592 | 0.006570357 | | | | | |
| | | | 2.95 | 0.37775 | 0.019711072 | | | | | |
| | | | 9.82 | 1.25917 | 0.065703574 | | | | | |
| | | | 29.47 | 3.77751 | 0.197110722 | | | | | |
| | | | 98.2 | 12.5917 | 0.65703574 | | | | | |
| | | | 294.72 | 37.7751 | 1.971107219 | | | | | |
| Trovafloxacin mesylate | 5μM[33] | 0.24 [61] | 0.1 | 0.12793 | 0.12 | | | | | |
| | | | 0.3 | 0.38380 | 0.36 | | | | | |
| | | | 1 | 1.27932 | 1.2 | | | | | |
| | | | 3 | 3.83795 | 3.6 | | | | | |
| | | | 10 | 12.7932 | 12 | | | | | |
| | | | 15 | 19.1898 | 18 | | | | | |
| | | | 30 | 38.3795 | 36 | | | | | |
| | | | 100 | 127.932 | 120 | | | | | |
| | | | 300 | 383.795 | 360 | | | | | |
| Ximelagatran | 0.45μM[53] | 1 | 0.1 | 0.045 | 0.045 | | | | | |
| | | | 0.3 | 0.135 | 0.135 | | | | | |
| | | | 1 | 0.45 | 0.45 | | | | | |
| | | | 3 | 1.35 | 1.35 | | | | | |
| | | | 10 | 4.5 | 4.5 | | | | | |
| | | | 30 | 13.5 | 13.5 | | | | | |
| | | | 100 | 45 | 45 | | | | | |
| | | | 300 | 135 | 135 | | | | | |
| Zileuton | 13.1μM[33] | 0.07 [62] | 0.1 | 0.1161 | 0.0917 | | | | | |
| | | | 0.3 | 0.3482 | 0.2751 | | | | | |
| | | | 1 | 1.1608 | 0.917 | | | | | |
| | | | 3 | 3.4823 | 2.751 | | | | | |
| | | | 10 | 11.608 | 9.17 | | | | | |
| | | | 30 | 34.823 | 27.51 | | | | | |
| | | | 100 | 116.08 | 91.7 | | | | | |
| | | | 300 | 348.23 | 275.1 | | | | | |

*FIRU values assumed same as Fialuridine

**A calculation error for Troglitazone led to slightly misaligned dosing concentrations such that the Multiplier $xC_{max}$ values aren't exact multiples of $C_{max}$

***Highest concentration limited by solubility

**Supplementary Table S2.** Drug information and dosing concentrations used in the investigation. Cycle-specific concentrations have been added for further clarity.

| Incremental DILI sensitivity | Tox FNR vs. Baseline | Clinical dev. success rate | Portfolio IRR | Portfolio eNPV ($m) | NPV of chip costs ($m) | IRR on chip investment | Portfolio NPV / NPV of base R&D costs | Steady state uplift in pre-tax profit (%) | Annualized industry-wide pre-tax profit uplift ($m) |
|---|---|---|---|---|---|---|---|---|---|
| 0.0% | 100.0% | 11.5% | 11.0% | -3.5 | -3.5 | -3.5% | -0.2% | -0.1% | (207) |
| 7.5% | 99.0% | 11.5% | 11.0% | 1.1 | -3.6 | 14.4% | 0.1% | 0.0% | 61 |
| 14.9% | 98.0% | 11.6% | 11.0% | 5.7 | -3.6 | 24.7% | 0.3% | 0.2% | 331 |
| 22.4% | 97.0% | 11.6% | 11.1% | 10.3 | -3.6 | 30.4% | 0.5% | 0.3% | 602 |
| 29.8% | 96.0% | 11.7% | 11.1% | 15.0 | -3.6 | 33.8% | 0.8% | 0.5% | 876 |
| 37.3% | 95.0% | 11.7% | 11.2% | 19.8 | -3.6 | 36.0% | 1.0% | 0.7% | 1,151 |
| 44.7% | 94.0% | 11.7% | 11.2% | 24.5 | -3.6 | 37.6% | 1.3% | 0.8% | 1,428 |
| 52.2% | 93.0% | 11.8% | 11.2% | 29.3 | -3.6 | 38.7% | 1.6% | 1.0% | 1,707 |
| 61.5% | 91.8% | 11.8% | 11.3% | 35.3 | -3.6 | 39.9% | 1.9% | 1.2% | 2,058 |
| 67.1% | 91.0% | 11.9% | 11.3% | 39.0 | -3.7 | 40.4% | 2.1% | 1.3% | 2,270 |
| 74.5% | 90.0% | 11.9% | 11.4% | 43.9 | -3.7 | 41.0% | 2.3% | 1.5% | 2,555 |
| 82.0% | 89.0% | 11.9% | 11.4% | 48.8 | -3.7 | 41.5% | 2.6% | 1.6% | 2,841 |
| **87.0%** | **88.3%** | **12.0%** | **11.4%** | **52.1** | **-3.7** | **41.8%** | **2.8%** | **1.8%** | **3,034** |
| 96.0% | 87.1% | 12.0% | 11.5% | 58.1 | -3.7 | 42.3% | 3.1% | 2.0% | 3,384 |

**Supplementary Table S3.** Portfolio value and industry profits increase with reductions in the false negative rate (FNR) of the preclinical toxicology assessment, which cause fewer toxic drugs to enter the clinic. The leftmost column tabulates the proportional improvement in DILI detection versus the base case, and the next column shows the improvement in the toxicology FNR relative to the model's base case. Since DILI is around 13% or tox failure in development, near total DILI sensitivity can reduce the FNR to around 87% of its base case value. Subsequent columns then show the clinical development success rate from entry into Phase I to launch, the internal rate of return (IRR) of the R&D portfolio, the NPV of the portfolio discounted to the time of drug launch, the capitalized cost of Liver-Chips used in assessing the portfolio (discounted to the time of launch), and the marginal IRR on the Liver-Chip investment. The remaining columns capture the value uplift due to FNR improvement as a percentage uplift of the portfolio's NPV relative to the baseline NPV of R&D, percentage uplift of steady state pre-tax profits, and the estimated increase in annual pre-tax profits for the small-molecule drug development industry. The row highlighted in dark gray relates to the improvement in FNR that may result from incorporating the Liver-Chip into DILI prediction workflows in accordance with the 87% sensitivity estimated by the present study. The rows highlighted in light gray correspond to the 95% confidence interval around this point estimate. The economic model behind these calculations is provided in full in the Supplementary Materials.
